# Integration of Motor Planning and Execution through Latent Structure Reorganization in the Posterior Parietal Cortex

**DOI:** 10.1101/2025.06.10.658832

**Authors:** Stefano Diomedi, Francesco Edoardo Vaccari, Kostas Hadjidimitrakis, Patrizia Fattori, Ivilin Peev Stoianov

**Affiliations:** Institute of Cognitive Sciences and Technologies, National Research Council of Italy, Padova 35137, Italy; Department of Biomedical and Neuromotor Sciences, University of Bologna, Bologna 40126, Italy

**Keywords:** Posterior Parietal Cortex, Motor control, Population Dynamics, Neural Subspaces, Dimensionality Reduction

## Abstract

The posterior parietal cortex (PPC) plays a central role in sensorimotor control, performing visuomotor transformations, supporting planning, and providing visual feedback. However, it is unknown how the neural populations in different PPC areas organize their activity during this process. It has been proposed that PPC activity reflects population-level dynamics rather than distinct subpopulations, raising the question of how the population flexibly reorganize between the two main phases of motor control, planning and execution. To address this question, we analyzed neural dynamics in three PPC areas (PE, PEc, V6A) in the context of a delayed reaching task, applying dimensionality reduction techniques. This approach allows identifying whether activity in each area is organized into independent or partially overlapping dynamics across task phases. We found evidence of area-specific population subspaces, distinct for movement planning and execution. Specifically, the analysis revealed that in PE, which is a predominantly somatomotor area, neural activity occupied nearly orthogonal subspaces between the two phases, suggesting independent dynamics for movement planning and execution. In contrast, in V6A and PEc, which are involved in visuomotor transformations, we identified both shared and exclusive subspaces, indicating a more flexible representation of motor information in these areas. Overall, our findings suggest that parietal circuits combine both separation and sharing of neural representation to support computations during the different movement stages, providing new insights into the role of the PPC in generating flexible motor behavior
.

**Significance Statement:** The posterior parietal cortex (PPC) plays a central role in sensorimotor control. How neural ensembles in the PPC transition from planning to executing goal-directed movements remains poorly understood. Using dimensionality reduction on macaque electrophysiological data from delayed reach tasks, we identify an anteroposterior gradient: posterior visuomotor areas exhibit both overlapping and segregated subspaces for planning and execution, whereas anterior somatomotor regions show predominantly orthogonal dynamics. This organizational gradient reflects persistent posterior encoding of action goals and posture, and more distinct anterior representations of body state across planning and execution. The topographical and functional shift from visuomotor to somatic coding aligns with hierarchical predictive coding accounts of sensorimotor control and advances our understanding of sensorimotor transformations in the PPC.

## Introduction

The parietal cortex plays a central role in motor control, acting as a key hub for integrating sensory information and coordinating motor responses based on predictions and feedback (1–5). During action planning, parietal neurons simultaneously encode multiple potential intentions across different modalities (6), regardless of the effector, based on possible contingencies and context, before selecting the final action (7, 8). During action execution, the parietal neural population is thought to implement a forward model that estimates the state of the effector using visual and proprioceptive feedback (9). The posterior parietal cortex (PPC) is anatomically divided by the intraparietal sulcus into a superior region—the superior parietal lobule (SPL)—and an inferior region—the inferior parietal lobule (IPL). The SPL, in particular, has been extensively investigated for its role in the planning and online control of reaching movements (10–14). Within the SPL, which shows a rostral-to-caudal gradient from somatosensory to visual dominance, three key areas are discussed here: PE, PEc and V6A ((15), Figure 1A). PE (Broadman area 5) is a somatic region involved in multi-joint coordination, proprioception, and posture (16). In addition, PE also encodes motor parameters such as the depth and direction of reaching movements (17), while visual modulations is absent in this area. In contrast, PEc and V6A (both parts of Broadman area 7) are visuomotor areas that exhibit both visual (18, 19) and proprioceptive (20, 21) modulation, in addition to encoding motor parameters related to voluntary eye and arm movements (22). Recently, a class of dimensionality reduction techniques has led to the emergence of the so-called neural state-space framework, a method for studying large-scale neural recordings (23–25). Using this framework, population activity has been found to explore only a limited, lowdimensional portion of the full neural space, termed a neural subspace. Instead of representing population activity as a point in the high-dimensional space defined by all recorded neurons, dimensionality reduction identifies a limited number of factors governing the evolution of population dynamics, leading to a representation in a lower-dimensional space (26).

**Fig. 1.**
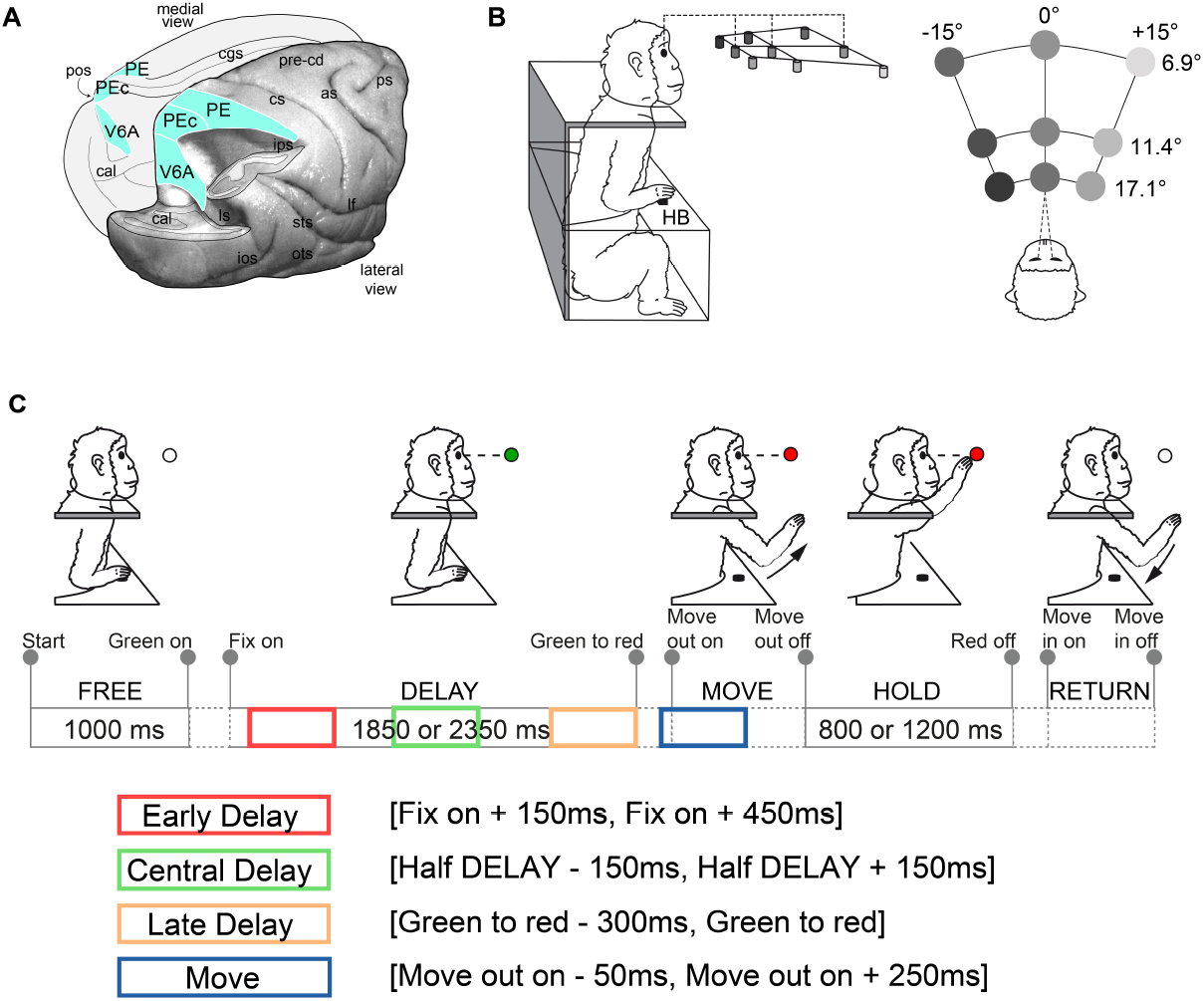
SPL areas and Behavioral Task. **A**. Posterolateral view of macaque brain. The right hemisphere is partially dissected at the level of the fundus of intraparietal, parieto-occipital, and lunate sulci to show the parieto-occipital sulcus. The areas PE, PEc and V6A are highlighted in yellow, with their boundaries outlined in white. Abbreviations: as, arcuate sulcus; cal, calcarine sulcus; cs, central sulcus; cgs, cingulate sulcus; ios, inferior occipital sulcus; ips, intraparietal sulcus; lf, lateral fissure; ls, lunate sulcus; ots, occipito-temporal sulcus; pos, parieto-occipital sulcus; pre-cd, pre-central dimple; ps, principal sulcus; sts, superior temporal sulcus. **B**. Lateral and top view of the experimental apparatus. Visual targets were arranged at the eye level on a single plane, varying in depth (ocular vergence: 17.1°, 11.4°, and 6.9°) and direction (ocular version: −15°, 0°, and +15°). **C**. Task sequence of foveated reaching task. The vignette illustrates the animal’s behavior using a color code. Grey circle indicate key time event (“Start”, “Green on”, “Fix on”, “Green to red”, “Move out on”, “Move out off”, “Red off”, “Move in on”, “Move in off”). Pairs of time events defined task epochs (“FREE”,”RT SACC”, “DELAY”, “RT MOVE OUT”, “MOVE”, “HOLD”, “RT MOVE IN”, “RETURN”). Task epochs that depends on animals’ behavior are shown as dashed rectangles, while those defined by task constraints (with fixed duration) are shown as solid rectangles, with duration labeled inside.

This approach helps to identify dynamic structures confined within specific generating subspaces thatare difficult to capture from single-neuron activity alone and enables the study of relationships between subspaces. For example, this framework has revealed that the motor cortex (M1) contains distinct and orthogonal neural subspaces dedicated to different stages of motor control (27). Elsayed et al. (28) found that neural activity associated with motor preparation operates independently of activity during movement execution, suggesting that M1 circuitry reorganizes to optimize performance across different phases of reaching tasks. Investigating M1 activity during both observed and executed movements, (29) found notable similarities in population covariation within a shared subspace. While the dynamics associated with observed and executed movements exhibit some overlap in the shared subspace, distinct and exclusive dynamics emerge in orthogonal subspaces dedicated to each condition. Notably, these dynamics do not result from the activation of a specific subpopulation of neurons but rather from the consistent temporal activation of a heterogeneous population.

Compared to the motor and premotor cortices, there is no clear evidence that PPC activity reorganizes to support its diverse sensorimotor functions. Single PPC neurons show a continuum of responses during goal-directed actions and have been categorized into sensory, motor, or sensorimotor cells (30, 31). However, (32), using Hidden Markov Models, showed that population activity in V6A, PEc, and PE transitions through distinct neural states during reaching, corresponding to preparation, execution, and post-movement stages (32, 33), similar to the findings in the motor cortices (34–36). Notably, these transitions engage the entire neural population rather than discrete subpopulations (32). This raises the hypothesis that PPC areas might flexibly perform multiple functions using a common neural substrate, a possibility that can be explored through neural subspace analysis. Such flexibility would require PPC activity to evolve through distinct subspaces depending on task demand.

Here, we investigate the relationship between the neural subspaces occupied by SPL populations during preparation and execution of reaching movements. We asked whether the preparatory and movement neural subspaces are orthogonal, indicating distinct processing stages, or whether overlapping dynamics occur within a common subspace, suggesting shared coding. We also examine whether exclusive subspaces emerge, dedicated specifically to preparation or movement.

We hypothesized that subspace organization would reflect the functional composition of each SPL area. In somatically dominated regions (PE; Brodmann Area 5), we expected neural dynamics to primarily reflect sensory coding, with little overlap between preparation and movement subspaces. In contrast, in more integrative regions containing a mix of sensory, sensorimotor, and motor neurons (PEc and V6A; Brodmann Area 7), we predicted both exclusive subspaces dedicated to preparation and execution, as well as a shared subspace supporting similar dynamics across these stages. Such a shared subspace would reflect a common neural substrate that integrates sensory input with motor planning and execution, facilitating a smooth transition between phases of motor control. By analyzing these relationships, we aim to uncover how SPL populations organize neural representations to support the transition from motor preparation to action execu-tion.

## Material and Methods

We reanalyzed publicly available data from our laboratory (37). The study was performed in accordance with the guidelines of the EU Directives (86/609/EEC; 2010/63/EU) and the Italian national law (D.L. 116-92, D.L. 26-2014) on the use of animals in scientific research. The protocols were approved by the Animal Welfare Body of the University of Bologna and by the Italian Ministry of Health. During the training and recording sessions, particular attention was paid to any behavioral and clinical signs of pain or distress.

### Experimental design and data preprocessing

#### Experimental procedure

Two male macaque monkeys (Macaca fascicularis), weighing 4.4 kg (*MonkeyS*) and 3.8 kg (*MonkeyF*), were used in this study. Extracellular single-cell activity was recorded from the anterior bank of the parieto-occipital sulcus (POs) and the adjacent caudal part of SPL. Multiple electrode penetrations were performed using a five-channel multielectrode recording system (Thomas Recording GmbH, Giessen, Germany). The electrode signals were amplified (with a gain of 10,000) and filtered (bandpass between 0.5 and 5 kHz). Action potentials in each channel were isolated using a waveform discriminator (Multi Spike Detector; Alpha Omega Engineering, Nazareth, Israel) and were sampled at 100 kHz. The quality of single-unit isolation was determined by visually inspecting spike waveforms and considering refractory periods in the inter-spike interval (ISI) histograms during spike sorting. Only well-isolated units with homogeneous waveforms and clear ISI histograms were considered.

The animal behavior was controlled using custom-made software implemented in the LabVIEW environment (National Instruments, Austin, TX)(38). Eye position signals were sampled using two cameras (one for each eye) of an infrared oculometer system (ISCAN, Woburn, MA) at 100 Hz. Although the vergence angle was not recorded online, it was reconstructed offline from the horizontal positions of the two eyes. A form of control for vergence was provided by the presence of electronic windows (one for each eye, 4° × 4° each) that controlled the frontoparallel gaze position, allowing us to offset the horizontal eye position signal for targets located in the same direction but at different depths.

The histological reconstructions of the electrode penetrations followed the procedures detailed in previous studies from our lab (18, 22).Area V6A was initially identified on functional grounds following the criteria described in (39) and later confirmed based on the cytoarchitectonic criteria reported in (40). The recording sites were assigned to area PEc according to the cytoarchitectonic criteria of (40) and (41). The PE recording site was determined following the cytoarchitectonic criteria of (41). Further details of the experimental methods are provided in (42).

#### Behavioral task

Electrophysiological signals were recorded while the monkeys performed an instructed delay-foveated reaching task in darkness (Figure 1B-C). The monkeys sat in a primate chair with their heads restrained and faced a horizontal panel at eye level. Nine light-emitting diodes (LEDs) on the panel served as fixation and reaching targets (Figure 1B). The LEDs were arranged in three rows: one central along the sagittal midline and two lateral rows at 15° to the left and right (Figure 1B). Each row contained three LEDs set at vergence angles of 17.1°, 11.4°, and 6.9°. Both monkeys had an interocular distance of 3.0 cm.

When performing the task, the monkeys used the limb opposite to the recording site while maintaining steady eye fixation. Each trial began when the monkey pressed a home button (HB, 2.5 cm in diameter, Figure 1B) located outside their visual field and 5 cm in front of their trunk. After 1 s, one of the nine LEDs lit up green, and the monkey had to fixate it while keeping the button pressed for 1700–2500 ms until the LED changed color from green to red, signaling the “Go” cue. The monkey then had 1 s to release the HB (movement onset) and start reaching for the target. The target touch was detected by a micro-switch at the base of the LED (movement end). After touching the target, the monkey had to hold it for 800–1200 ms. The task concluded with the LED turning off, prompting the monkeys to release the target and return to the initial position (HB) to receive a reward (Figure 1C). Only the trials in which the monkeys were rewarded were included in further analyses.

#### Data preprocessing

For each monkey (*MonkeyS, MonkeyF*), we considered three neural populations of single units one each for areas V6A, PEc and PE. The initial V6A, PEc, and PE populations were 157, 134, and 62 in *MonkeyS* and 128, 97 and 104 in *MonkeyF* respectively. From these initial populations, we retained neurons that exhibited a firing rate exceeding 3 spikes per seconds on avarage during main task epochs (FREE,DELAY,RT MOVE OUT, MOVE) and task-related neurons (one-way ANOVA, p-val<0.05, factor: epoch, 4 level, FREE, DELAY, RT MOVE OUT, MOVE see caption Figure 1C for task epochs notation). After applying these criteria, we continued the analysis with the following neural populations: *MonkeyS* 134 V6A, 116 PEc, and 48 PE and *MonkeyF* 99 V6A, 84 PEc, and 82 PE.

Due to the variability in trial and epoch duration, using a fixed temporal window results in a variable number of bins across trials. For this reason, we adopted a variable bin strategy. This approach allows us to create averaged trials and concatenate them into a matrix on wich we perform dimensionality reduction. For each animal, we calculated the epochs average duration across different conditions and brain areas. Subsequently, we determined the number of bins for those average epochs by dividing the average epoch duration by the desired bin size (10 ms). Next, for each trial, we can bin each epoch by selecting the desired number of bins, thereby achieving a bin duration very close to the original 10 ms (10.02 ± 0.0763 ms, mean ± standard deviation across monkeys).

After this procedure we computed firing rates by dividing the number of spikes occurring within the bin by the temporal length of the bin. We then smoothed the firing rates with a Gaussian kernel having standard deviation 80 ms. In the next step of preprocessing, we normalized firing rate following soft-normalization procedure (43), given the faring rate *f* :

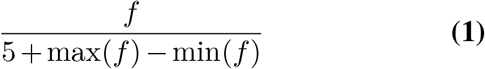

This method ensures that neurons exhibiting strong responses are constrained to a range of unity, while those with weaker responses are assigned firing rates of less than one. In the last preprocessing phase, the normalized firing rate has been mean-centered at each time: computing mean activity across all condition we subtract it for each condition response.

#### Epochs of interest and Neural Tensor

We were interested in the study of planning and movement phases. Regarding the planning phase, this takes place in the delay between the presentation of the target and the go cue. Given the high duration of this period, we have broken the delay phase into 3 sub-epochs of length about 300 ms each one. The first, Early Delay epoch, in order to avoid noise due to post saccadic response, started 150 ms after fixation onset and last for 300 ms. The second, Central Delay, was defined as the 300 ms time windows centered in the middle of the delay phase. The third, Late Delay was the last 300 ms in the delay phase before the go cue. Due to the pre-activation of muscle activity, which occurs approximately 50 ms before the actual start of movement, the Move epoch started 50 ms before the movement onset until 250 ms after it (see Figure 1C for a graphical scheme of the epochs selected). Each population analysis was performed on trial-averaging firing rates grouped into a 3D tensor defined in ℝ^*N×C×T*^ where *N* is the number of neurons, *C* is the number of target conditions and *T* is the number of time points. From a whole trial tensor, we cut the epoch of interest. In the next sessions we will explain methods for two general epochs called *EP*_*i*_ and *EP*_*j*_.

### Preliminary analysis

#### Cross correlations

For every epoch of interest *EP*_*i*_, we first computed the cross-correlation matrices between the activity of every couple of neurons. Tensors *EP*_*i*_ were reshaped into 2D tensors v^*N ×CT*^ and then the Person correlation coefficients between their rows were calculated to build cross-correlation matrices. We then investigated whether correlation structures were maintained or disrupted between two epochs. To that aim, we sorted the neurons according to the correlations in one of the epochs and then visualized the correlations in the other epoch using the same order. The order was established on the basis of hierarchical clustering of neural activity with the help of the *Linkage* function in Matlab 2023b. The cluster to which the neurons belonged was used to sort and the number of clusters was chosen empirically to enhance the emergence of structures. Moreover, the analysis was performed twice for every couple of epochs, so that each epoch was used once for sorting and twice for reordering. Finally, the relationship between every couple of correlation matrices was quantitatively assessed by correlating the two matrices using again the Pearson’s correlation.

#### Epoch-preference index

We assess the tuning strength 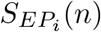 of a neuron *n* in an epoch *EP*_*i*_, as the ratio between the maximum of the neuron’s firing rate (across conditions) during the epoch and the average firing rate range of that neuron across all time. Then, given two epochs *EP*_*i*_ and *EP*_*j*_ we define an Epoch-preference index of a neuron as follows (28):

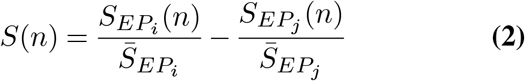

where 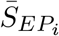 and 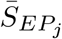 represent the mean tuning across all neurons. This index can take positive values if *EP*_*i*_ is preferred by the neuron, and negative values if *EP*_*j*_ is preferred. If the distribution of Epoch-preference indexes is bimodal (with two peaks, one on a negative value and one on a positive value), this means that there are two sub-populations of neurons, each one selective for a specific epoch. Bimodality was assessed using the Hartigan’s dip test.

### Identification of Neural Subspaces

#### Classic Principal Component Analysis and explained variance

We performed PCA separately for each epoch, treating the activity of each neuron as a variable of the input space. For each epoch, we used the first 10 principal components to compute the percentage of explained variance (called *self-variance*) and the normalized variance (see Normalized Variance section of *Materials and Methods*). This analysis identified a 10-dimensional *subspace* that maximizes the variance explained in a given epoch. Given a subspace and the corresponding transformation derived from the PCA computed over a given epoch, we then quantified how much variance in the neural activity of another epoch could be explained, which we call *cross-variance*.

***vOptimized Principal Component Analysis***

For detailed sub-spaces characterization, we used MATLAB toolbox *Manopt* (44) which was adapted according to (28) and (29). Here we briefly summarize the method; for further detail, we refer to the original publications.

### Orthogonal subspaces

The method searches two orthogonal subspaces. These are obtained by identifying a basis in the set of orthonormal matrices (Stiefel manifold) that maximizes an operator that expresses the sum of the normalized neural variance in the two epochs. The dimension of the Stiefel manifold was set as the number *d*_*i*_ of PCs (Principal Components) explaining 70% of the variability in *EP*_*i*_ plus the number *d*_*j*_ of PCs explaining 70% in the other epoch *EP*_*j*_.

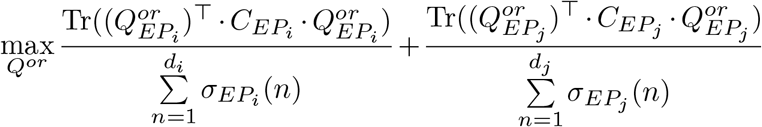

subject to

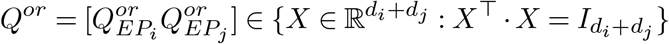

where 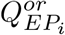 and 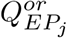 compose the basis of the two orthogonal subspaces. The notations 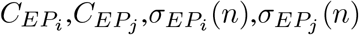 are explained in section Normalized Variance/Alignment Index.

### Exclusive subspaces

to find an Exclusive subspace we look for a base in the Stiefel manifold that maximizes a functional that expresses the normalized neural variance of one epoch with the additional constrains that the variance explained in the other epoch is less than a small threshold *v*:

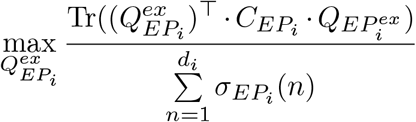

subject to

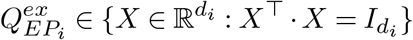

And

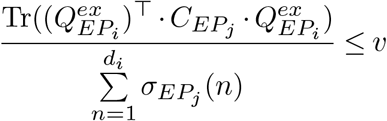

In Figure 5 we show results where *v* = 0.01.

**Shared subspaces**: the shared subspace was identified as the subspace that maximizes the neural variance explained in both epochs and that is orthogonal to the exclusive subspaces of both epochs, with dimension of the shared subspaces set at *d*_*sh*_ = min(*d*_*i*_, *d*_*j*_):

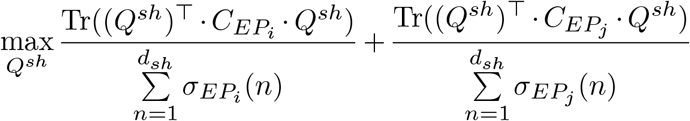

subject to:

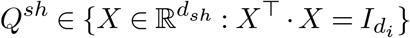

and

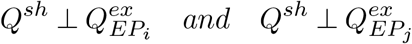

### Neural Subspace Analysis

#### Dimensionality Estimation

In the main analyses described in the Results section (Figure 4-5), the dimensionality of spaces was chosen following a common approach (28, 29, 45): a threshold of 70% was set and the dimensions needed to reach it were considered. To verify the sensitivity of the results to the dimensionality, we tested the variance explained in the subspaces optimized for various dimensions (Figure S7 and S11). However, for the linear decoding analysis, it was not possible to test high-dimensional spaces due to numerical instability in the regression model. Specifically, increasing the number of dimensions results in predictor matrices that are rank-deficient. This leads to numerical instability in matrix inversion, producing unreliable estimates (e.g., R^2^ values close to 1 both in real and shuffled data, and extremely low p-values). This issue is a known limitation of linear regression in high-dimensional settings and prevented us from including dimensions above a certain threshold. The number of dimensions tested was chosen based on simulations in our recent study (46), and was often consistent with the dimensionality estimated using a 70% variance threshold (Figure S8).

#### Normalized Variance

To assess whether two subspaces are overlapping, we use a metric called *normalized variance* (28) which quantifies the amount of variance shared between two subspaces *EP*_*i*_ and *EP*_*j*_ generated from different epochs. It calculates the overlap between two neural subspaces as the fraction of variance in one epoch explained by the other’s subspace. A value of 1 indicates complete overlap, that is, the subspace generated from *EP*_*i*_ fully explains the activity in *EP*_*j*_, whereas a value of 0 indicates that the subspaces are completely orthogonal. The index is computed as follows:

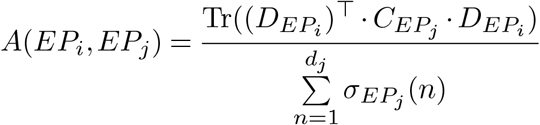

where 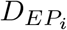 is the matrix defined by the *d*_*j*_ principal components selected in *EP*_*i*_, 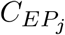 is the covariance of the 2D tensor ℝ^*N×CT*^ containing neural activity of *EP*_*j*_ activity, 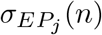 is the n-th singular value of 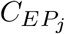 calculated.

To assess whether the observed overlap between subspaces from different epochs is greater than what would be expected by chance, we computed a baseline distribution of normalized variance. Specifically, we generated random subspaces by sampling orthonormal bases from the neural activity space, constrained by the empirical covariance structure of the data. This ensures that the synthetic subspaces have a dimensionality and variance structure comparable to those derived from real epochs. We then computed normalized variance between these random subspaces, repeating the procedure multiple times to build a null distribution. Comparing the empirical normalized variance to this null distribution allows us to determine whether the observed subspace overlap is statistically significant or could be explained by high-dimensional geometry alone. To assess significance, we performed a right-tailed test: for each pair of epochs, we computed a p-value as the proportion of random normalized variance that were greater than or equal to the empirical normalized variance.

#### Linear estimation

To asses whether the fluctuations in neural population activity projections during planning and execution reflect meaningful shared dynamics or random noise, we applied a linear estimation analysis between the two task epochs as in (28). First, we projected the full time-resolved neural population activity from each epoch (*EP*_*i*_ and *EP*_*j*_) onto their respective orthogonal subspaces obtained via optimized PCA. This yielded two time series of low-dimensional trajectories, one per epoch and condition. From these projected trajectories, we selected a single representative time slice per epoch, 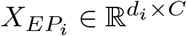 and 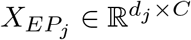. The slicing procedure was as follows: for each condition, we first located the time point within the epoch at which the absolute projection onto the first principal component (PC1) reached its maximum. We then took the median of these time points across all conditions and used that single value as the representative time slice for the epoch. We then fitted a linear model:

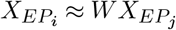

where 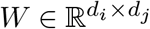 is the linear mapping. The system was solved using MATLAB’s backslash operator (\). Prediction accuracy was quantified with the coefficient of determination (*R*^2^). Furthermore, to assess statistical significance, we constructed a null distribution by shuffling the *C* dimension (columns of 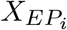 and 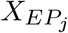) 10,000 times and re-fitting the model for each iteration.

## Results

We analyzed neural activity from two monkeys (*MonkeyF* and *MonkeyS*) performing an instructed delayed reaching task in darkness (for task details, see Methods and Figure 1 B-C). We compared population dynamics across three planning epochs (Early, Central, and Late Delay) and the movement epoch. Since the findings were consistent across all delay periods and animals, we focus here on the comparison between Early Delay and Move in *MonkeyF*. We choose to show Early Delay because it is functionally analogous to the preparatory period used in (28). A complete analysis of all delay epochs is provided in the Supplementary Material.

Our goal was to characterize the neural subspaces involved in movement planning and execution across three SPL areas, considering three possible scenarios: (i) that planning and execution subspaces are *coincident*, with population activity evolving along the same neural dimensions; (ii) that they are *orthogonal*, with planning and execution dynamics engaging distinct dimensions; and (iii) that *shared* and *exclusive* subspaces coexist, with population activity evolving partly along common neural dimensions (shared subspaces) and partly along phase-specific ones (exclusive subspaces).

### Cross-correlation analyses suggest independent epochs

As a preliminary step, we computed crosscorrelations between all pairs of neurons, separately for each cortical area and task epoch, to examine whether correlation structures emerging during planning persisted during movement execution (Figure 2A). We sorted the correlation matrices based on patterns of co-activation during Early Delay (see *Materials and Methods*). Notably, the structured patterns observed during Early Delay (visible as red and white regions) were substantially disrupted during movement, suggesting a reorganization of functional interactions across epochs (Figure 2A). To quantify this disruption, we computed the correlation between corresponding cross-correlation matrices across epochs. The resulting coefficients were low (*R*^2^: V6A, 0.09; PEc, 0.03; PE, 0.02; for *MonkeyS* see Figure S1), indicating that correlation structures during planning and execution were largely unrelated.

**Fig. 2.**
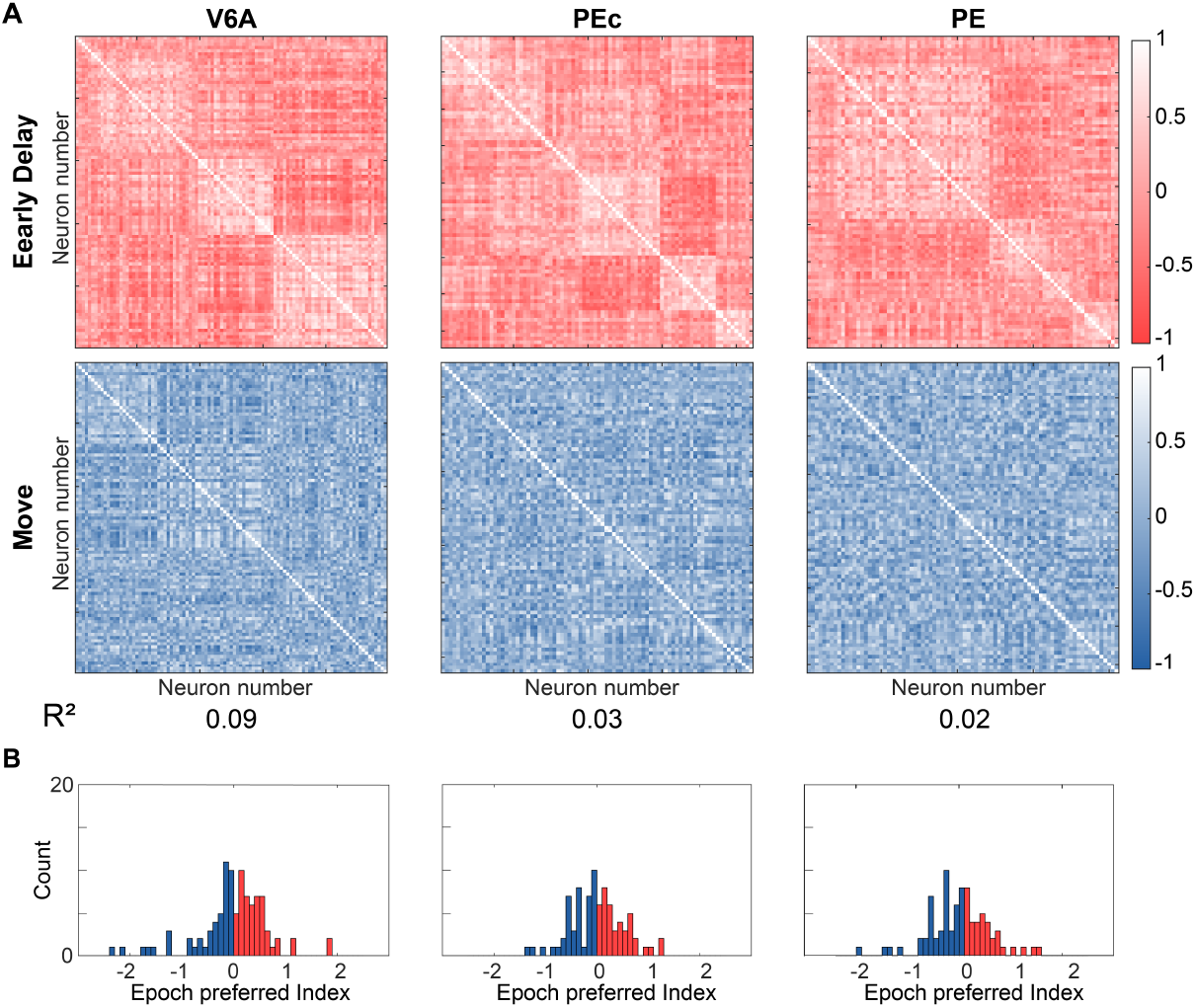
Cross-Correlation analysis. **A**. Cross-correlation matrices for each area and epoch (Early Delay: upper row, red; Movement: lower row, blue). Each matrix quantifies pairwise similarity in neuronal activity during the corresponding epoch (color scale). Neurons are ordered identically in both matrices per area. The *R*^2^ values at the bottom indicate the similarity between the Early Delay and Movement correlation structures. **B**. Epoch-preference for each neuron, grouped by area. Positive values indicate preference for Early Delay, negative for Movement. All distributions appeared unimodal, consistent with a continuous representation of activity across epochs (Table S1).

To ensure this result did not arise from distinct neuron sub-populations being active exclusively in one epoch, we assessed each neuron’s epoch preference using a Hartigan’s dip test on the distribution of an epoch-preference index (Figure 2B). A dip statistics close to zero indicates a unimodal distribution, suggesting a graded continuum of activity across epochs, while larger values imply bimodality potentially reflecting here two discrete subpopulations. The resulting dis-tributions were unimodal and centered around zero across all areas (all *B <* 0.03 and all *p* − *val >* 0.90; for *MonkeyS* see Table S1). This implies that the vast majority of neurons were active across both epochs, and that neurons selective for one epoch were rare. Similar results were obtained in the second animal and the other planning epochs (*SI* Figure S1-S2). These findings indicate that population activity patterns during planning and execution are relatively independent, challenging the idea that neural dynamics evolve within the same subspace across epochs. Based on this, we next asked whether the dynamics in the two task phases occupy orthogonal spaces.

### Latent Structure of Epochs

To probe the latent structure of neural representations during planning and movement, we performed principal component analysis (PCA) separately for each epoch. The activity of all neurons over time was used to construct a data matrix, from which PCA identified a low-dimensional representation that best captured the population dynamics. Specifically, we extracted the first ten principal components, defining a 10-dimensional latent subspace that maximized the variance explained within each epoch. The proportion of variance captured by this subspace in its own epoch was termed *self-variance*. To examine how neural representations generalize across epochs, we projected the population activity from one epoch onto the subspace derived from another and measured the explained variance-referred to as *cross-variance*. High cross-variance would suggest that the two epochs share similar latent structure, while low cross-variance suggests that planning and movement engage distinct, partially orthogonal subspaces. Thus, high self-variance paired with low cross-variance implies that each epoch is characterized by a unique latent representation.

Applying this approach to all SPL areas and both monkeys, we found that the 10-D Early Delay subspace captured 89%, 89%, and 87% of self-variance in V6A, PEc, and PE, respectively and explained 26%, 21%, and 16% of cross-variance (see Figure S4 for *MonkeyS*). Similarly, the Move subspace explained 89%, 88%, and 85% of self-variance and 27%, 18%, and 15% of cross-variance (see Figure S4 for *MonkeyS*). These findings suggest that although both epochs are internally consistent in their low-dimensional structure, their latent representations are only partially shared. These results are visualized in Figure 3A which breaks down the variance explained by each principle component. We further quantified the overlap – or alignment – between subspaces using a normalized cross-variance index, defined as the ratio of cross-variance to self-variance (see *Materials and Methods*). This measure ranges from zero (completely independent subspaces), to one (fully overlapping subspaces). For each area, the average normalized variance was relatively low V6A: 0.30, PEc: 0.23, PE: 0.18 (Figure 3B and see Table S2 for *MonkeyS*), indicating partial but significant alignment between the two task epochs.

**Fig. 3.**
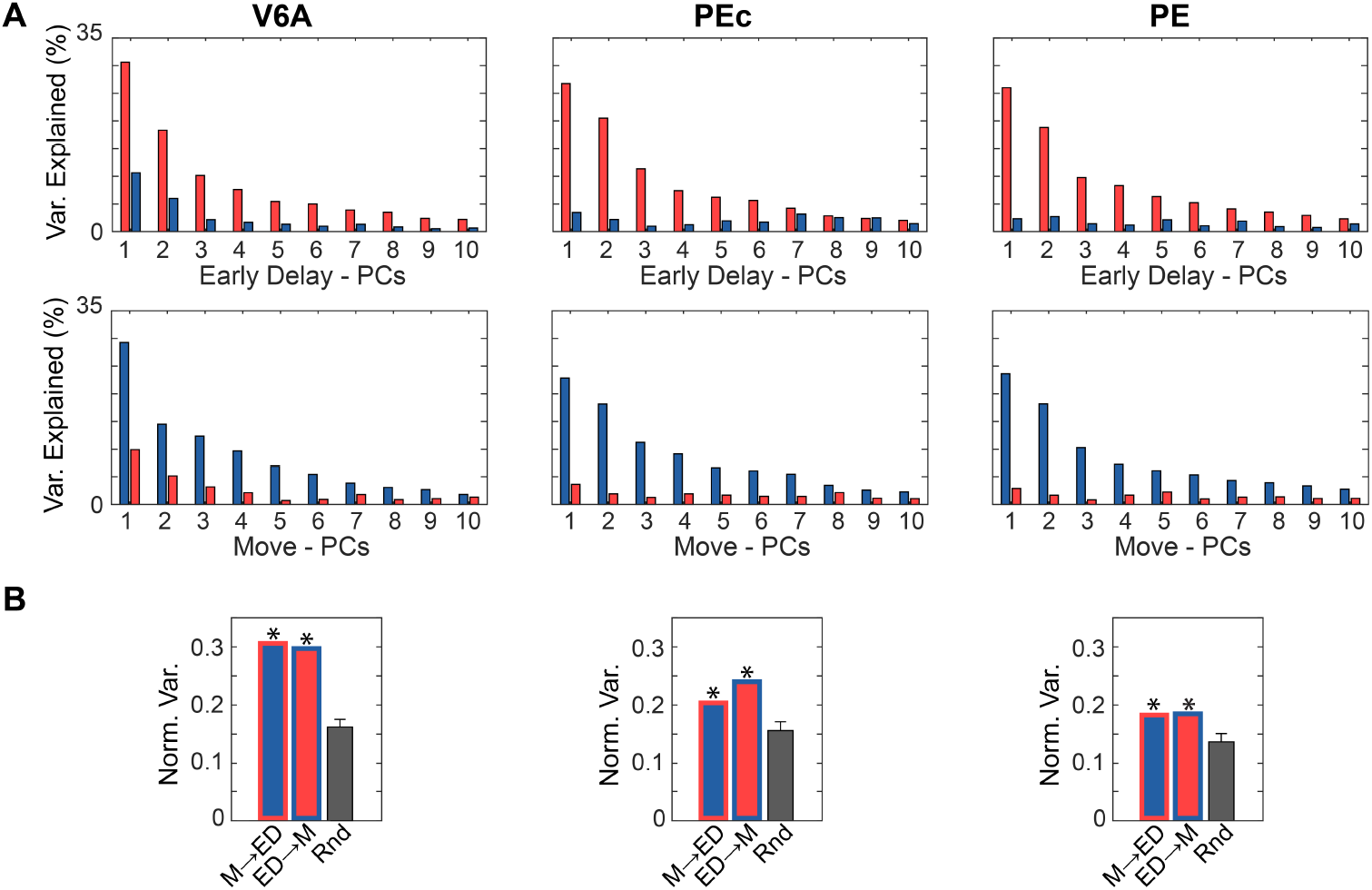
Principal Component Analysis. **A**. Percentage of variance explained by the top ten principle components (PCs). Top row: variance explained in the Early Delay (red bars) and Move (blue bars) epochs by PCs computed from the Early Delay epoch. Bottom row: variance explained by PCs computed from the Move epoch. **B**. Normalized variance captured by cross-projection of neural activity onto the 10-D subspaces. Bar outline color indicates the subspace, while fill color indicates the projected epoch (red: Early Delay; blue: Move). For example a blue bar with a red outline reflects the variance explained when Move activity is projected onto the Early Delay subspace (M*→*ED). Dark gray bars shown the mean and S.D. of normalized variance for 10.000 random subspaces (Rnd). Asterisks indicate normalized cross variance significantly greater than chance (all *p − val <* 0.001). Abbreviations: ED, Early Delay; M, Move; Rnd, Random.

Because high-dimensional subspaces can exhibit spurious overlap due to geometric constraints, we estimated the chance level of normalized cross-variance using a nullmodel. We generated random subspaces by projecting surrogate neural activity onto low-dimensional bases sampled from the empirical covariance structure (see *Materials and Methods*). This approach preserves some statistical features of the data while breaking the temporal and task-specific structure. The resulting null distribution (dark gray bars in Figure 3B) reflects the overlap expected by chance. In all areas, the observed subspace alignment was significantly higher than chance (*MonkeyF* all *p* − *val <* 0.001; see *Materials and Methods*), indicating that the overlap between planning and movement subspaces reflects a genuine structure rather than a statistical artifact.

Interestingly, in area PE, the average normalized cross-variance (mean of the blue and red bars in Figure 3B) was closer to the normalized chance level compared to other areas, suggesting a borderline result for this region. Similar patterns (most importantly, in terms of ratio) were observed for other delay epochs and the second monkey (see *SI* Figure S3-S4).

### Distinguishing Orthogonal and Non-Orthogonal Subspaces

The results thus far suggest that the latent structures do not fully overlap across epochs, nor that they are fully orthogonal. We investigated this issue further with an additional analysis that resolves a key methodological issue associated with a potential problem of interpretation. The PCAs were computed separately over different delays and epochs, which might lead to a fictive overlap that arises not from shared structure but from uncorrelated noise captured by the separate PCAs in a similar way. Thus, cross-variance might be due to noise that happened to be aligned in the two subspaces by chance and not because of a real functional relationship between the subspaces with actual behavioral meaning.

To address this issue, we applied an extension of PCA that seeks a common set of axes (or components) explaining as much variance as possible in both epochs while keeping the axes orthogonal (44); also see *Materials and Methods*. In this analysis, if the brain performs different computations during planning and execution, the corresponding activity would occupy separate (orthogonal) subspaces. If, instead, some amount of variance is shared between the two epochs, this would be captured through the cross-projections of activity onto the two orthogonal subspaces. To investigate whether some shared variance reflects meaningful neural signals or random noise, we used linear regression that verified whether the activity of one epoch could be predicted from the components of the other. If prediction was not possible, then any overlap would be likely due to noise rather than a functional link. Using this method, we identified an Early Delay Orth subspace that explained 93%, 98%, and 97%, and a Move Orth subspace that explained 93%, 97%, and 97% of the within-epoch normalized variance in V6A, PEc, and PE, respectively (see Table S3 for *MonkeyS*). Figure 4A shows the projections of neural activity onto the first latent dimension of each subspace, separately for each area and epoch. Neural dynamics were strongly modulated during their corresponding reference epoch, but also exhibited residual activity during the other epoch, which appeared as weak but consistent activations outside the preferred phase. This indicates that planning and execution dynamics cannot be cleanly separated. This observation was further supported by the normalized cross-variance, which reached approximately 10% (Figure 4B).

The critical question then was whether the observed 10% cross-variance could be attributed merely to noise—suggesting that the weaker activations reflected random fluctuations — or whether it indicated a meaningful relationship between the neural activity occurring during the two epochs. To address this, we used linear estimation to test whether activity patterns in the Early Delay Orth subspace (i.e., activity during the Early Delay epoch projected onto Early Delay Orth subspace) could predict activity in the Move Orth subspace (i.e., activity during the Move epoch projected onto Move Orth subspace), and vice versa. To assess whether the prediction reflected a meaningful relationship, we compared the fit values *R*^2^ with the fit of a chance-level distribution obtained by shuffling the independent variable 10, 000 times. Only predictions that significantly exceeded this baseline were considered indicative of a functional link between the two subspaces (see *Materials and Methods* for further details). The results of this analysis for *MonkeyF* are shown in Figure 4B. In epochs V6A and PEc, we observed significant predictive relationships between epochs, with fit values significantly greater than what would be expected by chance (*p − val <* 0.05, one-tailed test), suggesting a functional link between planning and execution dynamics. This was not the case in area PE, where the cross-predictive power did not exceed the chance level.

**Fig. 4.**
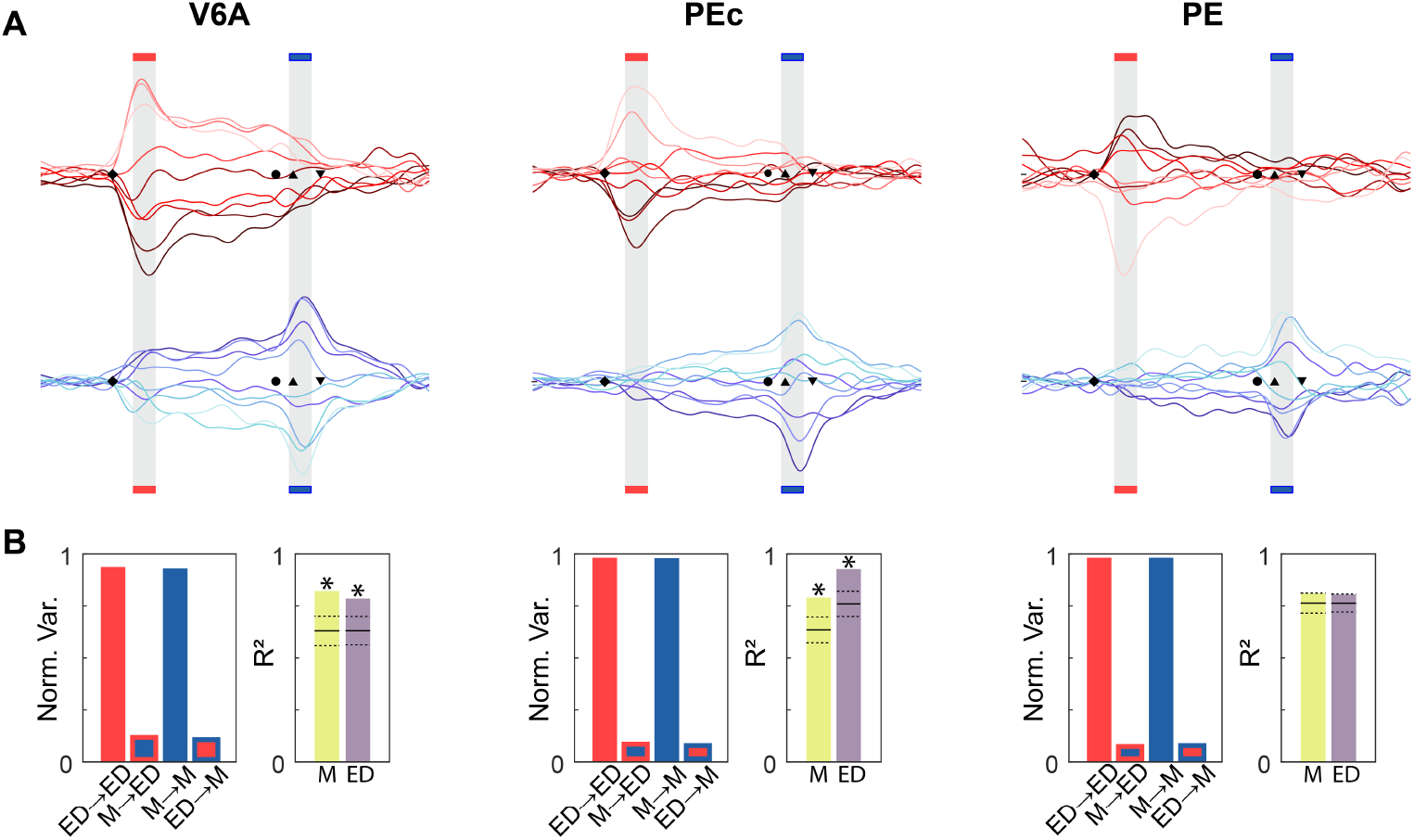
Orthogonal Subspaces. **A**. Projections of the neural population activity on the first principal component of the Early Delay Orthogonal Subspace (red traces) and onto the first principal component of the Move Orthogonal Subspace (blue traces). Lines of different color intensities represent different reach conditions, from left (darkest) to right (lightest).The black markers represent the main marker events of the task:*o*,green on;*?*, green to red;*6*, move out onset; *v*, move out offset. Compare sequence in Figure 1C. **B**. For each area, the left plot shows the Normalized Variance per epoch and subspace; outline and fill colors indicate subspace and epoch. In turn, the right plot shows the precision of the linear estimations. Yellow bar represents the *R*^2^ fit of the estimation of Move dynamics in Move subspace from Early Delay projection in the Early Delay subspace, while the Pink bar shows the opposite. Asterisks shows significant difference relative to a null distribution (*p − val <* 0.05, one-tailed test). Null distribution was obtained by shuffling the independent variable of the fitting, which was repeated 10.000 times. The mean fit of the random shuffling is represented by horizontal continuous line and the standard deviation with a horizontal dashed line. Abbreviations: ED, Early Delay; M, Move.

In summary, the orthogonality analysis partially confirmed the findings obtained with classic PCA: computations in V6A and PEc are not fully orthogonal between planning and movement epochs, whereas in PE, population dynamics appeared largely segregated across the two. A similar trend was observed for *MonkeyS*, also comparing the Central Delay epoch and Late Delay epoch (Figure S5–S6). Based on these results, we concluded the analysis for PE and proceeded to further investigate shared and exclusive subspaces in V6A and PEc.

### Distinguishing Exclusive and Shared Subspaces

Results so far demonstrate that the computations for planning and execution in V6A and PEc are not orthogonal, indicating the presence of a shared subspace. We investigated the structure of the subspaces further, considering two possible scenarios: (i) that one subspace A is entirely contained within another subspace B (A ⊂ B), resulting in a shared structure and a single exclusive component (A = shared, B \ A = exclusive); (ii) that the two subspaces intersect partially, each maintaining a portion of exclusive variance beyond their shared components (A *∩* B = shared, A *\* shared = A-exclusive, B \shared = B-exclusive). To distinguish between these scenarios, we applied another version of PCA (see *Materials and Methods*) capable of identifying both shared and exclusive subspaces.

**Fig. 5.**
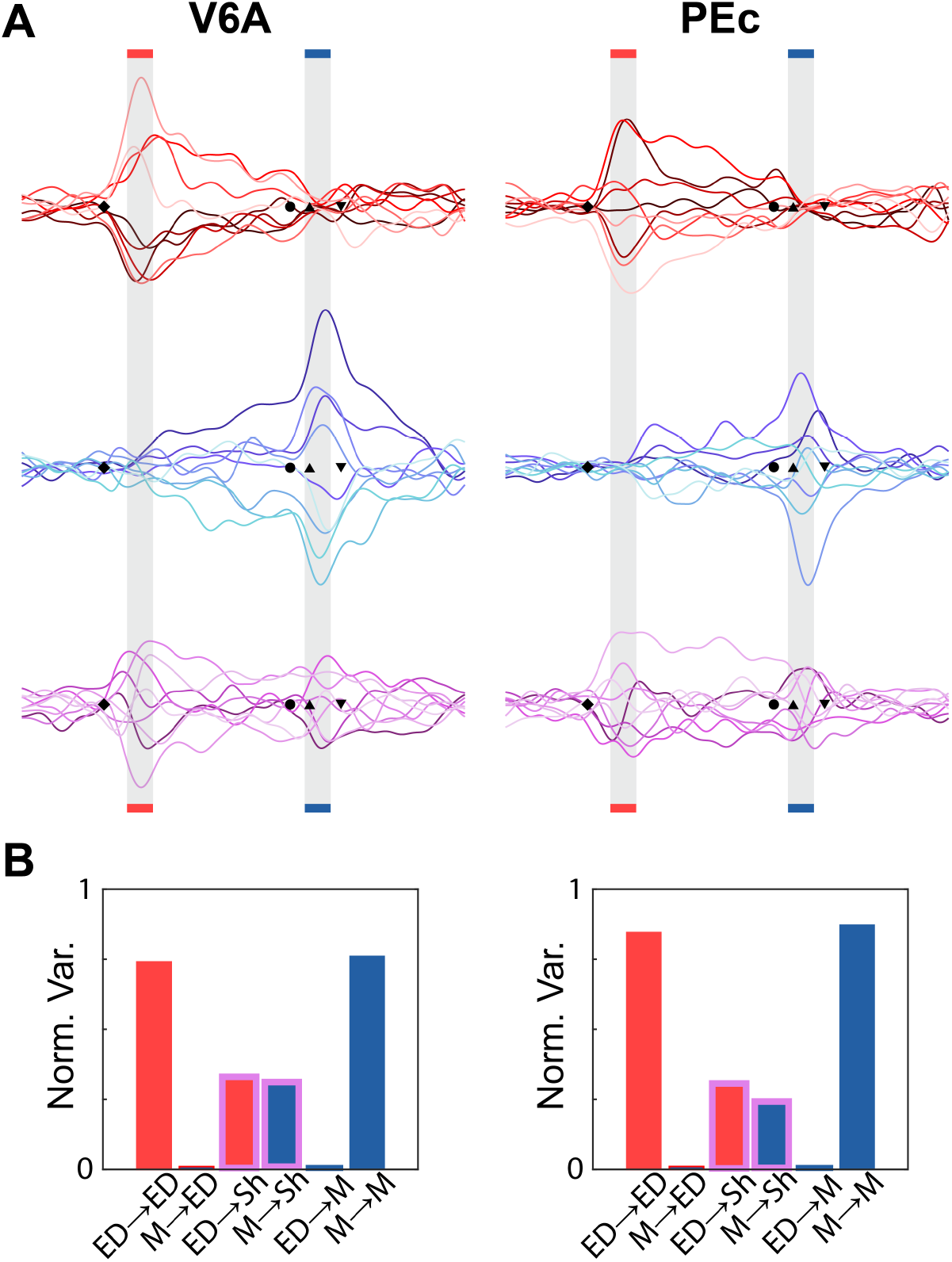
Exclusive and Shared subspaces. **A**. Projections of the neural population activity on the first principal component of the Early Delay Exclusive Subspace (red traces), onto the first principal component of the Move Exclusive Subspace (blue traces) and onto the first principal component of the Shared subspaces (magenta traces). Lines of different color intensities represent different reach conditions, from left (darkest) to right (lightest) targets. **B**. Normalized Variance relative to Exclusive and Shared subspaces: outline and fill color indicate subspace and epoch (red, ED Ex; blue, M Ex; magenta, Shared). Abbreviations: ED, Early Delay; M, Move; Sh, Shared.

As shown in Figure 5B, the results of the analysis supported the second hypothesis. First, in both V6A and PEc, we identified exclusive subspaces in both planning (Early Delay) and execution (Move). In V6A (PEc), the Exclusive Early Delay subspace captured 74% (84%) of the normalized self-variance across animals, while the Exclusive Move subspace captured 76% (87%). In turn, the shared subspace explained 33% (30%) of the normalized planning variance and 30% (24%) of the execution variance in V6A (PEc), (see Table S4 for *MonkeyS*; compare also Figure S9-S10). Most importantly, when projecting the population activity onto the first principal component of each exclusive subspace, we observed clear segregation: dynamics associated with movement execution were only active during the Move epoch and absent or minimal during planning; conversely, preparatory activity was present only in the Early Delay epoch and vanished during movement.

## Discussion

We investigated neural population dynamics across three regions of the posterior parietal cortex during the planning and execution phases of a reaching task, using a state-space framework. Our goal was to determine whether neural activity evolves within fully overlapping, fully orthogonal subspaces, or partially overlapping subspaces across these distinct behavioral epochs. Correlation analysis and classical PCA revealed that planning and execution engage largely distinct subspaces: each epoch’s subspace primarily explained its own variance, and only minimally accounted for activity in the other epoch (Figures 2, 3). This finding rules out a fully overlapping representation of planning and movement and points instead to distinct activation patterns across epochs. However, the degree of subspace orthogonality varied across regions as revealed by analysis with an optimized PCA. In V6A and PEc, we observed partial overlap, indicating shared dynamics between planning and execution. By contrast, PE showed stronger orthogonality, suggesting largely independent subspaces for each subspace. Linear estimation analysis reinforced these results: In V6A and PEc, both shared and exclusive (epoch-specific) subspaces were identified, consistent with partially overlapping population codes(Figure 5). In contrast, neural dynamics in PE were predominantly exclusive to each epoch (Figure 4B). Together, our findings suggest a gradient of population-level organization along the caudal-to-rostral axis of the SPL – from partially overlapping latent structures in V6A and PEc to more segregated representations in PE. This shift in latent dynamics may reflect a functional transition from sensorimotor integration in caudal PPC to more specialized motor computations in rostral PPC.

### The lack of full overlap suggests mixed selectivity and dynamic sensorimotor encoding

The partial separation of neural subspaces across task epochs has been documented in motor cortices (27, 28, 47), where distinct cognitive or motor processes give rise to unique neural population dynamics that occupy distinct or partially overlapping subspaces (43). In our study, the incomplete separation between planning and execution is consistent with this view. Evidence for discrete neural states associated with different task phases has also emerged from studies using Hidden Markov Models, which identified planning-and execution-specific states in V6A, PEc, and PE (32, 48). In motor cortex, spatial tuning properties have been shown to shift between planning and movement, suggesting dynamic and context-dependent encoding (28, 49). Similarly, in parietal areas, spatial preferences evolve during reaching – from stronger direction selectivity during planning to movement extent during execution (50). This dissociation could contribute to the partially orthogonal subspaces we observed. Support for this interpretation also comes from human PPC, where orthogonal coding of shoulder and hand movements has been reported (30), paralleling findings of orthogonal planning and execution axes in motor areas (51). Such signal segregation may reflect functional specialization, for instance, between direction- and extent-related variables, or between sensory and motor transformations. Furthermore, parietal areas such as V6A, PEc, and PE play key roles in visuomotor transformations across different reference frames during reach tasks, a process associated with distinct encoding formats across task epochs (52, 53) – potentially contributing to the emergence of independent subspaces.

#### Independent computations during planning and execution in somatomotor area PE

The distinct subspaces we observed in PE during planning and execution likely reflect specialized computations unique to each task phase. Planning is dominated by anticipatory processes such as target localization, motor intention, and posture estimation (52, 54–56), while execution engages motor coordination, sensory feedback processing, and movement fine-tuning (57). This functional separation may support computational efficiency by partitioning cognitive and motor operations into largely segregated neural circuits. Our analysis revealed that PE exhibits purely orthogonal dynamics between epochs. As a low-level association area, PE integrates somatic information with motor signals to support posture and limb movement (15, 16, 58, 59), and it contains a detailed topographic map of the upper limbs (60). It receives strong input from primary somatosensory cortex (58), likely generating distinct sensory states – static during planning and dynamic during execution – that result in divergent population activity patterns. Because PE encodes the current limb state via sensory feedback, rather than abstract movement goals, the population activity during planning and execution may occupy orthogonal subspaces. Notably, PE is also connected to M1 (58) and projects to the spinal cord (61). In motor cortices, preparatory activity is known to occupy a “null-space” that is orthogonal to movement-driving dimensions (27). Similarly, the orthogonal subspaces observed in PE may reflect a strategy to minimize interference between internal estimations of body states and outgoing motor commands or incoming sensory feedback. During planning, PE may encode predicted proprioceptive states, which are then updated through sensory prediction errors during execution – key for accurate and adaptive motor control (1, 62, 63).

### Independent and shared computations within visuo-motor areas V6A and PEc

Unlike PE, areas V6A and PEc are primarily visuomotor, integrating visual and somatosensory information during both planning and execution of goal-directed actions (10, 64). These regions encode spatial object locations and support coordinated eye and hand movements (15). During planning, they combine visuospatial cues to build motor intentions; during execution, they process sensory feedback to adjust ongoing actions (3, 65, 66). Exclusive subspaces during planning likely reflect goal prediction (52), build-up of motor intentions (67), and expected outcomes of action (4, 68). In contrast, exclusive subspaces during execution capture real-time adjustments based on real-time feedback – consistent with predictive and active inference theories of motor control (67–70).

Furthermore, unlike PE, both V6A and PEc exhibited not only exclusive but also shared subspaces between planning and execution. This suggests partial reuse of neural circuitry across task phases, and more generally, within the broader process of motor control. Shared subspaces may enhance computational efficiency by maintaining a common representational format for related computations. Indeed, similar shared dynamics have been observed across different tasks, suggesting generalized processing mechanisms (29, 71, 72). Within a single task, shared dynamics may reflect behavioral features that remain constant across epochs, for example, sustained fixation. Thus, the presence of shared subspaces in V6A and PEc may stem from their modulation by eye position signals (16, 52, 73, 74), which are weak or absent in PE. More importantly, both areas maintain stable encoding of the reach target across task phases (48, 75), likely contributing to the emergence of shared latent codes that span planning and execution and most likely represent motor intentions ().

### Exclusive and shared subspaces may respectively reflect the encoding of body and environment states

The PPC integrates forward predictions and sensory feedback to estimate the current state of the body and the environment (55, 66, 76). Recent evidence suggests that these computations are spatially organized along the rostrocaudal axis: rostral PPC predominantly process body-related signals, while caudal PPC represents environmental information (2).

In our experiment, the external environment remained constant – the animal performed the task in a fixed workspace – whereas the body state changed markedly between the static delay period and dynamic movement execution. These differences were reflected in the structure of the latent subspaces. Exclusive subspaces, particularly pronounced in rostral area PE, may capture body-related representations that differ across task phases, from static to dynamic. In contrast, shared subspaces – primarily found in caudal areas V6A and PEc – likely encode stable aspects of the environment that remain constant across task phases. This interpretation is consistent with prior findings showing that caudal PPC areas are involved in stable environment encoding (55, 77), whereas rostral areas contribute more to dynamic body state estimation (9, 78). Together, these results suggest that the PPC maintains distinct but interactive representations of the body and environment, integrating them dynamically to guide behavior. The coexistance of exclusive and shared subspaces may reflect this integration, supporting predictive control of action under stable and changing task constraints (68).

### The functional role of V6A, PEc, and PE: a synthesis

The present study reveals different organization of planning and execution subspaces in medial PPC: V6A and PEc exhibit both exclusive and shared subspaces, while PE shows largely orthogonal dynamics. This pattern aligns with known anatomical and functional gradients along the PPC’s rostrocaudal axis. The anterior area PE primarily processes somatosensory input (58, 79), whereas the more posterior V6A and PEc receive stronger visual inputs (80–83). Notably, these areas also belong to different Brodmann regions, with V6A and PEc to area 7 and PE to area 5 (15).

The clear subspace segregation in PE is consistent with its role in encoding arm posture and movement, and its strong connectivity with M1 (17, 58, 84). This result parallels findings in M1, where orthogonal but linked subspaces segregate planning and execution dynamics (28), enabling preparatory activity to prime motor computations (27, 43). In PE, orthogonal subspaces may separate static sensory states from dynamic motor outputs, reducing interference between preparation and execution.

In contrast, V6A and PEc – less influenced by somatosensory delays (3, 66) – likely integrate visuomotor signals within a shared subspace, enabling a smoother transition from planning to execution. Unlike the rapid subspace configurations in the frontal cortex (27, 85), the shared subspace reconfiguration in PPC may reflect a gradual sensorimotor transformation process. While M1 maintains stable sensorimotor tuning throughout task execution, PPC exhibits a progression from sensory to motor coding as the task unfolds (86). The shared subspaces in V6A and PEc may serve to pre-align sensorimotor information, preparing downstream motor areas for accurate execution (28, 85).

### Limitations and future perspectives

Our findings have several methodological limitations. The animals performed a highly constrained reaching task in complete darkness with fixed target positions—conditions that enhance neural interpretability but may limit generalization to naturalistic behaviors. Additionally, by using trial-averaged neural activity and temporal smoothing, we were able to leverage single-unit recordings and reduce noise, enabling the identification of population-level subspace dynamics. However, this approach may obscure trial-by-trial variability and fast transient dynamics critical for understanding motor control and error correction.

Future research should explore how subspace geometry varies across individual trials, especially regarding behavioral variability, movement errors, and learning. It would also be valuable to investigate how PPC subspaces interact with premotor and motor cortices, to determine whether shared subspaces are coordinated across regions or arise from locally organized dynamics.

## Conclusions

Together, our findings reveal a principled organization of latent neural dynamics along the posterior-to-anterior axis of the medial PPC. V6A and PEc exhibit a balance of exclusive and shared subspaces, consistent with their role in integrating visual and somatosensory information to support predictive and flexible visuomotor transformations. In contrast, the predominantly orthogonal structure in PE reflects its specialization for proprioceptive and motor processing, likely optimized for the separation of planning and execution-related signals. These distinctions mirror known anatomical and functional gradients across the PPC, suggesting that subspace organization reflects the functional priorities of each region – ranging from predictive coding and sensory integration in caudal PPC to motor-specific encoding in rostral areas. Our study thus highlights how population-level neural dynamics are adapted to support distinct computational roles across the parietal reach pathway, contributing to the seamless transformation of intention into action.

## Data Availability

All neural and behavioural data used for this work are publicly available at https://doi.gin.g-node.org/10.12751/gnode.7q2dbp/. For an extensive description of the datasets see (42).

## Author contributions

Stefano Diomedi: Conceptualization; Data curation; Formal analysis; Investigation; Methodology; Validation; Visualization; Writing – original draft; Writing – review & editing. Francesco Edoardo Vaccari: Visualization; Writing – review & editing. Kostas Hadjidimitrakis: Supervision; Writing – review & editing. Patrizia Fattori: Resources; Writing – review & editing. Ivilin Peev Stoianov: Supervision; Funding acquisition; Writing – review & editing.

## ACKNOWLEDGEMENTS

We would like to thank Prof. Margherita Porcelli for her helpful comments. Generative AI was used to correct typographical errors and edit language for clarity.

## Bibliography

1. Matteo Priorelli and Ivilin Peev Stoianov. Flexible Intentions: An Active Inference Theory. Frontiers in Computational Neuroscience, 17:1 – 41, 2023. doi: 10.3389/fncom.2023.1128694.

2. W. Pieter Medendorp and Tobias Heed. State estimation in posterior parietal cortex: Distinct poles of environmental and bodily states. Progress in Neurobiology, 183:101691, December 2019. ISSN 03010082. doi: 10.1016/j.pneurobio.2019.101691.

3. Patrizia Fattori, Rossella Breveglieri, Annalisa Bosco, Michela Gamberini, and Claudio Galletti. Vision for Prehension in the Medial Parietal Cortex. Cerebral Cortex, 27(February): 1149–1163, 2017. ISSN 1047-3211. doi: 10.1093/cercor/bhv302.

4. Rick A. Adams, Stewart Shipp, and Karl J. Friston. Predictions not commands: active inference in the motor system. Brain Structure and Function, 218(3):611–643, May 2013. ISSN 1863-2653, 1863-2661. doi: 10.1007/s00429-012-0475-5.

5. Stephen H. Scott. Optimal feedback control and the neural basis of volitional motor control. Nature Reviews Neuroscience, 5(7):532–545, July 2004. ISSN 1471-003X, 1471-0048. doi: 10.1038/nrn1427.

6. Shenbing Kuang, Pierre Morel, and Alexander Gail. Planning Movements in Visual and Physical Space in Monkey Posterior Parietal Cortex. Cerebral Cortex, page bhu312, January 2015. ISSN 1047-3211, 1460-2199. doi: 10.1093/cercor/bhu312.

7. He Cui. From Intention to Action: Hierarchical Sensorimotor Transformation in the Posterior Parietal Cortex. eneuro, 1(1):ENEURO.0017–14.2014, November 2014. ISSN 2373-2822. doi: 10.1523/ENEURO.0017-14.2014.

8. Christian Klaes, Stephanie Westendorff, Shubhodeep Chakrabarti, and Alexander Gail. Choosing Goals, Not Rules: Deciding among Rule-Based Action Plans. Neuron, 70(3):536–548, May 2011. ISSN 08966273. doi: 10.1016/j.neuron.2011.02.053.

9. Grant H. Mulliken, Sam Musallam, and Richard A. Andersen. Forward estimation of movement state in posterior parietal cortex. Proceedings of the National Academy of Sciences, 105(24):8170–8177, June 2008. ISSN 0027-8424, 1091-6490. doi: 10.1073/pnas.0802602105.

10. Patrizia Fattori, Marina De Vitis, Matteo Filippini, Francesco Edoardo Vaccari, Stefano Diomedi, Michela Gamberini, and Claudio Galletti. Visual sensitivity at the service of action control in posterior parietal cortex. Frontiers in Physiology, 15:1408010, May 2024. ISSN 1664-042X. doi: 10.3389/fphys.2024.1408010.

11. Rossella Breveglieri, Sara Borgomaneri, Stefano Diomedi, Alessia Tessari, Claudio Galletti, and Patrizia Fattori. A Short Route for Reach Planning between Human V6A and the Motor Cortex. The Journal of Neuroscience, 43(12):2116–2125, March 2023. ISSN 0270-6474, 1529-2401. doi: 10.1523/JNEUROSCI.1609-22.2022.

12. Claudio Galletti, Dieter F. Kutz, Michela Gamberini, Rossella Breveglieri, and Patrizia Fattori. Role of the medial parieto-occipital cortex in the control of reaching and grasping movements. Experimental Brain Research, 153(2):158–170, November 2003. ISSN 0014-4819, 1432-1106. doi: 10.1007/s00221-003-1589-z.

13. Aaron P. Batista and Richard A. Andersen. The Parietal Reach Region Codes the Next Planned Movement in a Sequential Reach Task. Journal of Neurophysiology, 85(2):539– 544, February 2001. ISSN 0022-3077, 1522-1598. doi: 10.1152/jn.2001.85.2.539.

14. S. Ferraina, M. R. Garasto, A. Battaglia-Mayer, P. Ferraresi, P. B. Johnson, F. Lacquaniti, and R. Carniniti. Visual Control of Hand-reaching Movement: Activity in Parietal Area 7m. European Journal of Neuroscience, 9(5):1090–1095, May 1997. ISSN 0953-816X, 1460-9568. doi: 10.1111/j.1460-9568.1997.tb01460.x.

15. Michela Gamberini, Lauretta Passarelli, Patrizia Fattori, and Claudio Galletti. Structural connectivity and functional properties of the macaque superior parietal lobule. Brain Structure and Function, 225(4):1349–1367, May 2020. ISSN 1863-2653, 1863-2661. doi: 10.1007/s00429-019-01976-9.

16. Marina De Vitis, Rossella Breveglieri, Konstantinos Hadjidimitrakis, Wim Vanduffel, Claudio Galletti, and Patrizia Fattori. The neglected medial part of macaque area PE: segregated processing of reach depth and direction. Brain Structure and Function, 224(7):2537–2557, September 2019. ISSN 1863-2653, 1863-2661. doi: 10.1007/s00429-019-01923-8.

17. Kostas Hadjidimitrakis, Marina De Vitis, Masoud Ghodrati, Matteo Filippini, and Patrizia Fattori. Anterior-posterior gradient in the integrated processing of forelimb movement direction and distance in macaque parietal cortex. Cell Reports, 41(6):111608, November 2022. ISSN 22111247. doi: 10.1016/j.celrep.2022.111608.

18. Michela Gamberini, Claudio Galletti, Annalisa Bosco, Rossella Breveglieri, and Patrizia Fattori. Is the Medial Posterior Parietal Area V6A a Single Functional Area? The Journal of Neuroscience, 31(13):5145–5157, March 2011. ISSN 0270-6474, 1529-2401. doi: 10.1523/JNEUROSCI.5489-10.2011.

19. A. Battaglia-Mayer. Eye-Hand Coordination during Reaching. II. An Analysis of the Relationships between Visuomanual Signals in Parietal Cortex and Parieto-frontal Association Projections. Cerebral Cortex, 11(6):528–544, June 2001. ISSN 14602199. doi: 10.1093/cercor/11.6.528.

20. R. Breveglieri, C. Galletti, S. Monaco, and P. Fattori. Visual, Somatosensory, and Bimodal Activities in the Macaque Parietal Area PEc. Cerebral Cortex, 18(4):806–816, April 2008. ISSN 1047-3211, 1460-2199. doi: 10.1093/cercor/bhm127.

21. Rossella Breveglieri, Dieter F. Kutz, Patrizia Fattori, Michela Gamberini, and Claudio Galletti. Somatosensory cells in the parieto-occipital area V6A of the macaque:. NeuroReport, 13(16):2113–2116, November 2002. ISSN 0959-4965. doi: 10.1097/00001756-200211150-00024.

22. Michela Gamberini, Giulia Dal Bò, Rossella Breveglieri, Sofia Briganti, Lauretta Passarelli, Patrizia Fattori, and Claudio Galletti. Sensory properties of the caudal aspect of the macaque’s superior parietal lobule. Brain Structure and Function, December 2018. ISSN 1863-2653, 1863-2661. doi: 10.1007/s00429-017-1593-x.

23. Arthur Pellegrino, Heike Stein, and N. Alex Cayco-Gajic. Dimensionality reduction beyond neural subspaces with slice tensor component analysis. Nature Neuroscience, 27(6):1199– 1210, June 2024. ISSN 1097-6256, 1546-1726. doi: 10.1038/s41593-024-01626-2.

24. David Thura, Jean-François Cabana, Albert Feghaly, and Paul Cisek. Integrated neural dynamics of sensorimotor decisions and actions. PLOS Biology, 20(12):e3001861, December 2022. ISSN 1545-7885. doi: 10.1371/journal.pbio.3001861.

25. John P Cunningham and Byron M Yu. Dimensionality reduction for large-scale neural recordings. Nature Neuroscience, 17(11):1500–1509, November 2014. ISSN 1097-6256, 1546-1726. doi: 10.1038/nn.3776.

26. Juan A. Gallego, Matthew G. Perich, Lee E. Miller, and Sara A. Solla. Neural Manifolds for the Control of Movement. Neuron, 94(5):978–984, June 2017. ISSN 08966273. doi: 10.1016/j.neuron.2017.05.025.

27. Matthew T Kaufman, Mark M Churchland, Stephen I Ryu, and Krishna V Shenoy. Cortical activity in the null space: permitting preparation without movement. Nature Neuroscience, 17(3):440–448, March 2014. ISSN 1097-6256, 1546-1726. doi: 10.1038/nn.3643.

28. Gamaleldin F Elsayed, Antonio H Lara, Matthew T Kaufman, Mark M Churchland, and John P Cunningham. Reorganization between preparatory and movement population responses in motor cortex. Nat. Commun., 7(1):13239, October 2016.

29. Xiyuan Jiang, Hemant Saggar, Stephen I Ryu, Krishna V Shenoy, and Jonathan C Kao. Structure in neural activity during observed and executed movements is shared at the neural population level, not in single neurons. Cell Rep., 32(6):108006, August 2020.

30. Carey Y. Zhang, Tyson Aflalo, Boris Revechkis, Emily R. Rosario, Debra Ouellette, Nader Pouratian, and Richard A. Andersen. Partially Mixed Selectivity in Human Posterior Parietal Association Cortex. Neuron, 95(3):697–708.e4, August 2017. ISSN 08966273. doi: 10.1016/j.neuron.2017.06.040.

31. Hideo Sakata, Yoshio Takaoka, Atsushi Kawarasaki, and Hidetoshi Shibutani. Somatosensory properties of neurons in the superior parietal cortex (area 5) of the rhesus monkey. Brain Research, 64:85–102, December 1973. ISSN 00068993. doi: 10.1016/0006-8993(73)90172-8.

32. S. Diomedi, F.E. Vaccari, C. Galletti, K. Hadjidimitrakis, and P. Fattori. Motor-like neural dynamics in two parietal areas during arm reaching. Progress in Neurobiology, 205:102116, October 2021. ISSN 03010082. doi: 10.1016/j.pneurobio.2021.102116.

33. Stefano Diomedi. Neural states in parietal areas during arm reaching. University of Bologna, 2021. doi: 10.48676/unibo/amsdottorato/9991.

34. Kevin A. Mazurek and Marc H. Schieber. Mirror neurons precede non-mirror neurons during action execution. Journal of Neurophysiology, 122(6):2630–2635, December 2019. ISSN 0022-3077, 1522-1598. doi: 10.1152/jn.00653.2019.

35. Kevin A. Mazurek, Adam G. Rouse, and Marc H. Schieber. Mirror Neuron Populations Represent Sequences of Behavioral Epochs During Both Execution and Observation. The Journal of Neuroscience, 38(18):4441–4455, May 2018. ISSN 0270-6474, 1529-2401. doi: 10.1523/JNEUROSCI.3481-17.2018.

36. Caleb Kemere, Gopal Santhanam, Byron M. Yu, Afsheen Afshar, Stephen I. Ryu, Teresa H. Meng, and Krishna V. Shenoy. Detecting Neural-State Transitions Using Hidden Markov Models for Motor Cortical Prostheses. Journal of Neurophysiology, 100(4):2441–2452, October 2008. ISSN 0022-3077, 1522-1598. doi: 10.1152/jn.00924.2007.

37. Stefano Diomedi, Francesco E. Vaccari, Michela Gamberini, Marina De Vitis, Matteo Filippini, and Patrizia Fattori. Single-cell recordings from three cortical parietal areas during an instructed-delay reaching task. G-Node, March 2024. doi: 10.12751/G-NODE.7Q2DBP.

38. D. F. Kutz, N. Marzocchi, P. Fattori, S. Cavalcanti, and C. Galletti. Real-Time Supervisor System Based on Trinary Logic to Control Experiments With Behaving Animals and Humans. Journal of Neurophysiology, 93(6):3674–3686, June 2005. ISSN 0022-3077, 1522-1598. doi: 10.1152/jn.01292.2004.

39. Claudio Galletti, Patrizia Fattori, Dieter F. Kutz, and Michela Gamberini. Brain location and visual topography of cortical area V6A in the macaque monkey. European Journal of Neuroscience, 11(2):575–582, February 1999. ISSN 0953-816X, 1460-9568. doi: 10.1046/j.1460-9568.1999.00467.x.

40. Giuseppe Luppino, Suliann Ben Hamed, Michela Gamberini, Massimo Matelli, and Claudio Galletti. Occipital (V6) and parietal (V6A) areas in the anterior wall of the parieto-occipital sulcus of the macaque: a cytoarchitectonic study. European Journal of Neuroscience, 21 (11):3056–3076, June 2005. ISSN 0953-816X, 1460-9568. doi: 10.1111/j.1460-9568.2005.04149.x.

41. Deepak N. Pandya and Benjamin Seltzer. Intrinsic connections and architectonics of posterior parietal cortex in the rhesus monkey. Journal of Comparative Neurology, 204(2):196–210, January 1982. ISSN 0021-9967, 1096-9861. doi: 10.1002/cne.902040208.

42. S. Diomedi, F. E. Vaccari, M. Gamberini, M. De Vitis, M. Filippini, and P. Fattori. Neurophysiological recordings from parietal areas of macaque brain during an instructed-delay reaching task. Scientific Data, 11(1):624, June 2024. ISSN 2052-4463. doi: 10.1038/s41597-024-03479-7.

43. Mark M. Churchland, John P. Cunningham, Matthew T. Kaufman, Justin D. Foster, Paul Nuyujukian, Stephen I. Ryu, and Krishna V. Shenoy. Neural population dynamics during reaching. Nature, 487(7405):51–56, July 2012. ISSN 0028-0836, 1476-4687. doi: 10.1038/nature11129.

44. Nicolas Boumal, Bamdev Mishra, P.-A. Absil, and Rodolphe Sepulchre. Manopt, a matlab toolbox for optimization on manifolds. J. Mach. Learn. Res., 15(1):1455–1459, jan 2014. ISSN 1532-4435.

45. Antonio H Lara, Gamaleldin F Elsayed, Andrew J Zimnik, John P Cunningham, and Mark M Churchland. Conservation of preparatory neural events in monkey motor cortex regardless of how movement is initiated. eLife, 7:e31826, August 2018. ISSN 2050-084X. doi: 10.7554/eLife.31826.

46. Francesco Edoardo Vaccari, Stefano Diomedi, Edoardo Bettazzi, Matteo Filippini, Marina De Vitis, Kostas Hadjidimitrakis, and Patrizia Fattori. More or less latent variables when analysing neurophysiological data? this is the question. Bioarxiv, 2024.

47. Cheng Tang, Roger Herikstad, Aishwarya Parthasarathy, Camilo Libedinsky, and ShihCheng Yen. Minimally dependent activity subspaces for working memory and motor preparation in the lateral prefrontal cortex. eLife, 9:e58154, September 2020. ISSN 2050-084X. doi: 10.7554/eLife.58154.

48. Francesco Edoardo Vaccari, Stefano Diomedi, Marina De Vitis, Matteo Filippini, and Patrizia Fattori. Similar neural states, but dissimilar decoding patterns for motor control in parietal cortex. Network Neuroscience, 8(2):486–516, July 2024. ISSN 2472-1751. doi: 10.1162/netn_a_00364.

49. Nicholas G. Hatsopoulos, Qingqing Xu, and Yali Amit. Encoding of Movement Fragments in the Motor Cortex. The Journal of Neuroscience, 27(19):5105–5114, May 2007. ISSN 0270-6474, 1529-2401. doi: 10.1523/JNEUROSCI.3570-06.2007.

50. Kostas Hadjidimitrakis, Giulia Dal Bo’, Rossella Breveglieri, Claudio Galletti, and Patrizia Fattori. Overlapping representations for reach depth and direction in caudal superior parietal lobule of macaques. Journal of Neurophysiology, 114(4):2340–2352, October 2015. ISSN 0022-3077, 1522-1598. doi: 10.1152/jn.00486.2015.

51. Mark M. Churchland, John P. Cunningham, Matthew T. Kaufman, Stephen I. Ryu, and Krishna V. Shenoy. Cortical Preparatory Activity: Representation of Movement or First Cog in a Dynamical Machine? Neuron, 68(3):387–400, November 2010. ISSN 08966273. doi: 10.1016/j.neuron.2010.09.015.

52. Antonio Roberto Buonfiglio, Stefano Diomedi, Matteo Filippini, Patrizia Fattori, and Ivilin Peev Stoianov. Dynamic predictive spatial encoding of motor intentions in area v6a of the posterior parietal cortex. bioRxiv, 2025. doi: 10.1101/2025.02.06.636800.

53. Kostas Hadjidimitrakis, Masoud Ghodrati, Rossella Breveglieri, Marcello G. P. Rosa, and Patrizia Fattori. Neural coding of action in three dimensions: Task- and time-invariant reference frames for visuospatial and motor-related activity in parietal area V6A. Journal of Comparative Neurology, 528(17):3108–3122, December 2020. ISSN 0021-9967, 1096-9861. doi: 10.1002/cne.24889.

54. Jean-Jacques Orban de Xivry and Robert Hardwick. A control policy can be adapted to task demands during both motor execution and motor planning. bioRxiv, 2023. doi: 10.1101/2023.10.16.562495.

55. Yuhui Li, Yong Wang, and He Cui. Posterior parietal cortex predicts upcoming movement in dynamic sensorimotor control. Proceedings of the National Academy of Sciences, 119(13):e2118903119, March 2022. ISSN 0027-8424, 1091-6490. doi: 10.1073/pnas.2118903119.

56. Aaron L. Wong, Adrian M. Haith, and John W. Krakauer. Motor Planning.pdf. The Neuroscientist, 21(4), 2015. doi: 10.1177/107385841454148.

57. Daqi Dong and Stan Franklin. Sensory motor system: Modeling the process of action execution. Cognitive Science, 36, 2014.

58. Sophia Bakola, Lauretta Passarelli, Michela Gamberini, Patrizia Fattori, and Claudio Galletti. Cortical Connectivity Suggests a Role in Limb Coordination for Macaque Area PE of the Superior Parietal Cortex. The Journal of Neuroscience, 33(15):6648–6658, April 2013. ISSN 0270-6474, 1529-2401. doi: 10.1523/JNEUROSCI.4685-12.2013.

59. Stephen H. Scott, Lauren E. Sergio, and John F. Kalaska. Reaching Movements With Similar Hand Paths but Different Arm Orientations. II. Activity of Individual Cells in Dorsal Premotor Cortex and Parietal Area 5. Journal of Neurophysiology, 78(5):2413–2426, November 1997. ISSN 0022-3077, 1522-1598. doi: 10.1152/jn.1997.78.5.2413.

60. Adele M. H. Seelke, Jeffrey J. Padberg, Elizabeth Disbrow, Shawn M. Purnell, Gregg Recanzone, and Leah Krubitzer. Topographic Maps within Brodmann’s Area 5 of Macaque Monkeys. Cerebral Cortex, 22(8):1834–1850, August 2012. ISSN 1460-2199, 1047-3211. doi: 10.1093/cercor/bhr257.

61. Jean-Alban Rathelot, Richard P. Dum, and Peter L. Strick. Posterior parietal cortex contains a command apparatus for hand movements. Proceedings of the National Academy of Sciences, 114(16):4255–4260, April 2017. ISSN 0027-8424, 1091-6490. doi: 10.1073/pnas.1608132114.

62. Andrew B. Schwartz. Movement: How the Brain Communicates with the World. Cell, 164 (6):1122–1135, March 2016. ISSN 00928674. doi: 10.1016/j.cell.2016.02.038.

63. Wen Fang, Junru Li, Guangyao Qi, Shenghao Li, Mariano Sigman, and Liping Wang. Statistical inference of body representation in the macaque brain. Proceedings of the National Academy of Sciences, 116(40):20151–20157, October 2019. ISSN 0027-8424, 1091-6490. doi: 10.1073/pnas.1902334116.

64. Rossella Breveglieri, Riccardo Brandolani, Stefano Diomedi, Markus Lappe, Claudio Galletti, and Patrizia Fattori. Role of the medial posterior parietal cortex in orchestrating attention and reaching. Journal of Neuroscience, 45(1), 2025. ISSN 0270-6474. doi: 10.1523/JNEUROSCI.0659-24.2024.

65. Sohrab Saberi-Moghadam, Simone Ferrari-Toniolo, Stefano Ferraina, Roberto Caminiti, and Alexandra Battaglia-Mayer. Modulation of Neural Variability in Premotor, Motor, and Posterior Parietal Cortex during Change of Motor Intention. The Journal of Neuroscience, 36(16):4614–4623, April 2016. ISSN 0270-6474, 1529-2401. doi: 10.1523/JNEUROSCI.3300-15.2016.

66. Richard A. Andersen, Spencer Kellis, Christian Klaes, and Tyson Aflalo. Toward More Versatile and Intuitive Cortical Brain–Machine Interfaces. Current Biology, 24(18):R885–R897, September 2014. ISSN 09609822. doi: 10.1016/j.cub.2014.07.068.

67. Matteo Priorelli, Giovanni Pezzulo, and Ivilin Peev Stoianov. Deep kinematic inference affords efficient and scalable control of bodily movements. Proceedings of the National Academy of Sciences of the United States of America, 120:1–9, 2023. doi: 10.1073/pnas.2309058120.

68. Matteo Priorelli, Ivilin Peev Stoianov, and Giovanni Pezzulo. Embodied decisions as active inference. bioRxiv, 2025. doi: 10.1101/2024.05.28.596181.

69. Karl Friston. The free-energy principle: a unified brain theory? Nature Reviews Neuroscience, 11(2):127–138, February 2010. ISSN 1471-003X, 1471-0048. doi: 10.1038/nrn2787.

70. Matteo Priorelli, Federico Maggiore, Antonella Maselli, Francesco Donnarumma, Domenico Maisto, Francesco Mannella, Ivilin Peev Stoianov, and Giovanni Pezzulo. Modeling motor control in continuous time active inference: A survey. IEEE Transactions on Cognitive and Developmental Systems, 16(2):485–500, 2024. doi: 10.1109/TCDS.2023.3338491.

71. Giovanni Pezzulo, Francesco Donnarumma, Simone Ferrari-Toniolo, Paul Cisek, and Alexandra Battaglia-Mayer. Shared population-level dynamics in monkey premotor cortex during solo action, joint action and action observation. Progress in Neurobiology, 210:102214, March 2022. ISSN 03010082. doi: 10.1016/j.pneurobio.2021.102214.

72. Davide Albertini, Marco Lanzilotto, Monica Maranesi, and Luca Bonini. Largely shared neural codes for biological and nonbiological observed movements but not for executed actions in monkey premotor areas. Journal of Neurophysiology, 126(3):906–912, September 2021. ISSN 0022-3077, 1522-1598. doi: 10.1152/jn.00296.2021.

73. Rossella Breveglieri, Kostas Hadjidimitrakis, Annalisa Bosco, Silvio P. Sabatini, Claudio Galletti, and Patrizia Fattori. Eye Position Encoding in Three-Dimensional Space: Integration of Version and Vergence Signals in the Medial Posterior Parietal Cortex. The Journal of Neuroscience, 32(1):159–169, January 2012. ISSN 0270-6474, 1529-2401. doi: 10.1523/JNEUROSCI.4028-11.2012.

74. M. Raffi, A. Ballabeni, M.G. Maioli, and S. Squatrito. Neuronal responses in macaque area PEc to saccades and eye position. Neuroscience, 156(3):413–424, October 2008. ISSN 03064522. doi: 10.1016/j.neuroscience.2008.08.018.

75. A. Roberto Buonfiglio. Normalization of spatial coding for planning in the posterior parietal cortex. unknown, 2024.

76. He Cui. Forward Prediction in the Posterior Parietal Cortex and Dynamic Brain-Machine Interface. Frontiers in Integrative Neuroscience, 10, October 2016. ISSN 1662-5145. doi: 10.3389/fnint.2016.00035.

77. Yoh Inoue, Hongwei Mao, Steven B. Suway, Josue Orellana, and Andrew B. Schwartz. Decoding arm speed during reaching. Nature Communications, 9(1):5243, December 2018. ISSN 2041-1723. doi: 10.1038/s41467-018-07647-3.

78. Sebastian Dowiasch, Gunnar Blohm, and Frank Bremmer. Neural correlate of spatial (mis-)localization during smooth eye movements. European Journal of Neuroscience, 44(2):1846–1855, 2016. doi: 10.1111/ejn.13276.

79. Jeffrey Padberg, Dylan F. Cooke, Christina M. Cerkevich, Jon H. Kaas, and Leah Krubitzer. Cortical connections of area 2 and posterior parietal area 5 in macaque monkeys. Journal of Comparative Neurology, 527(3):718–737, February 2019. ISSN 0021-9967, 1096-9861. doi: 10.1002/cne.24453.

80. Lauretta Passarelli, Marcello G. P. Rosa, Michela Gamberini, Sophia Bakola, Kathleen J. Burman, Patrizia Fattori, and Claudio Galletti. Cortical Connections of Area V6Av in the Macaque: A Visual-Input Node to the Eye/Hand Coordination System. The Journal of Neuroscience, 31(5):1790–1801, February 2011. ISSN 0270-6474, 1529-2401. doi: 10.1523/JNEUROSCI.4784-10.2011.

81. S. Bakola, M. Gamberini, L. Passarelli, P. Fattori, and C. Galletti. Cortical Connections of Parietal Field PEc in the Macaque: Linking Vision and Somatic Sensation for the Control of Limb Action. Cerebral Cortex, 20(11):2592–2604, November 2010. ISSN 1047-3211, 1460-2199. doi: 10.1093/cercor/bhq007.

82. Michela Gamberini, Lauretta Passarelli, Patrizia Fattori, Mino Zucchelli, Sophia Bakola, Giuseppe Luppino, and Claudio Galletti. Cortical connections of the visuomotor parietooccipital area V6Ad of the macaque monkey. Journal of Comparative Neurology, 513(6):622–642, April 2009. ISSN 0021-9967, 1096-9861. doi: 10.1002/cne.21980.

83. Rossella Breveglieri, Claudio Galletti, Michela Gamberini, Lauretta Passarelli, and Patrizia Fattori. Somatosensory Cells in Area PEc of Macaque Posterior Parietal Cortex. The Journal of Neuroscience, 26(14):3679–3684, April 2006. ISSN 0270-6474, 1529-2401. doi: 10.1523/JNEUROSCI.4637-05.2006.

84. Stefano Ferraina, Emiliano Brunamonti, Maria Assunta Giusti, Stefania Costa, Aldo Genovesio, and Roberto Caminiti. Reaching in Depth: Hand Position Dominates over Binocular Eye Position in the Rostral Superior Parietal Lobule. The Journal of Neuroscience, 29(37):11461–11470, September 2009. ISSN 0270-6474, 1529-2401. doi: 10.1523/JNEUROSCI.1305-09.2009.

85. David Sussillo, Mark M Churchland, Matthew T Kaufman, and Krishna V Shenoy. A neural network that finds a naturalistic solution for the production of muscle activity. Nature Neuroscience, 18(7):1025–1033, July 2015. ISSN 1097-6256, 1546-1726. doi: 10.1038/nn.4042.

86. Cong Zheng, Qifan Wang, and He Cui. Continuous sensorimotor transformation enhances robustness of neural dynamics to perturbation in macaque motor cortex, November 2024.

