## Supplementary Materials for "Integration of Motor Planning and Execution through Latent Structure Reorganization in the Posterior Parietal Cortex"

DRAFT

| | *MonkeyF* | | *MonkeyS* | |
|---|---|---|---|---|
| | **B** | **p-value** | **B** | **p-value** |
| V6A | 0.03 | 0.94 | 0.03 | 0.50 |
| PEc | 0.03 | 0.94 | 0.01 | 1.00 |
| PE | 0.03 | 0.92 | 0.03 | 1.00 |

**Table S1. Hartigan's dip test for *MonkeyF* and *MonkeyS***
Hartigan's dip test statistics (B) and corresponding p-values for the distribution of epoch-preference indices, computed separately for each monkey and SPL area (V6A, PEc, PE). Low B values and high p-values indicate unimodal distributions, suggesting a continuous represenation of neural activity across epochs.

DRAFT

| | ***MonkeyF*** | | | | | ***MonkeyS*** | | | | |
|---|---|---|---|---|---|---|---|---|---|---|
| | Early Delay | | Move | | | Early Delay | | Move | | |
| | self var.(%) | cross var.(%) | self var.(%) | cross var.(%) | Norm. Var. | self var.(%) | cross var. (%) | self var.(%) | cross var. (%) | Norm. Var |
| V6A | 89 | 26 | 89 | 27 | 0.30 | 88 | 21 | 89 | 20 | 0.22 |
| PEc | 89 | 21 | 88 | 18 | 0.23 | 88 | 18 | 88 | 18 | 0.20 |
| PE | 87 | 16 | 85 | 15 | 0.18 | 84 | 26 | 86 | 28 | 0.24 |

**Table S2. Classic PCA subspace analysis for *MonkeyF* and *MonkeyS***
For each brain area (V6A, PEc, PE) and each epoch (Early Delay and Move ), the table reports the percentage of explained self and cross variance and the normalized variance between the two subspaces.

DRAFT

| | ***MonkeyF*** | | | | ***MonkeyS*** | | | |
|---|---|---|---|---|---|---|---|---|
| | Early Delay Orth | | Move Orth | | Early Delay Orth | | Move Orth | |
| | dim | self var. (%) | dim | self var. (%) | dim | self var. (%) | dim | self var. (%) |
| V6A | 5 | 93 | 5 | 93 | 5 | 95 | 5 | 94 |
| PEc | 5 | 98 | 6 | 97 | 5 | 96 | 5 | 95 |
| PE | 6 | 97 | 6 | 97 | 7 | 95 | 6 | 97 |

**Table S3. Orthogonal (Orth) subspace analysis for *MonkeyF* and *MonkeyS***
For each brain area (V6A, PEc, PE) and each epoch (Early Delay and Move ), the table reports the dimensionality of the subspace (dim) and the percentage of explained variance accounted for by that subspace (self var. %). Results highlight the low-dimensional structure capturing the majority of neural variance during both planning and movement execution phases.

DRAFT

| | *MonkeyF* | | | | | | *MonkeyS* | | | | | |
|---|---|---|---|---|---|---|---|---|---|---|---|---|
| | Early Delay Ex | | Move Ex | | Shared | | Early Delay Ex | | Move Ex | | Shared | |
| | dim | var. (%) | dim | var. (%) | dim | var. (%) | dim | var. (%) | dim | var. (%) | dim | var. (%) |
| V6A | 5 | 74 | 5 | 76 | 5 | 33 ED 30 M | 5 | 83 | 5 | 84 | 5 | 20 ED 20 M |
| PEc | 5 | 84 | 6 | 87 | 5 | 30 ED 24 M | 5 | 85 | 5 | 87 | 5 | 19 ED 21 M |

**Table S4. Shared and Exclusive (Ex) subspace analysis for *MonkeyF* and *MonkeyS***
Shared and Exclusive (Ex) subspace analysis for *MonkeyF* and *MonkeyS*, including the Early Delay and Move epochs. For each brain area (V6A, PEc), the table reports the dimensionality (dim) and the percentage of explained variance (var. %) captured by each epoch-specific subspace and by the shared subspace. Variance values for the shared subspace are specified separately for the Early Delay and Movement epochs (ED, M).

DRAFT

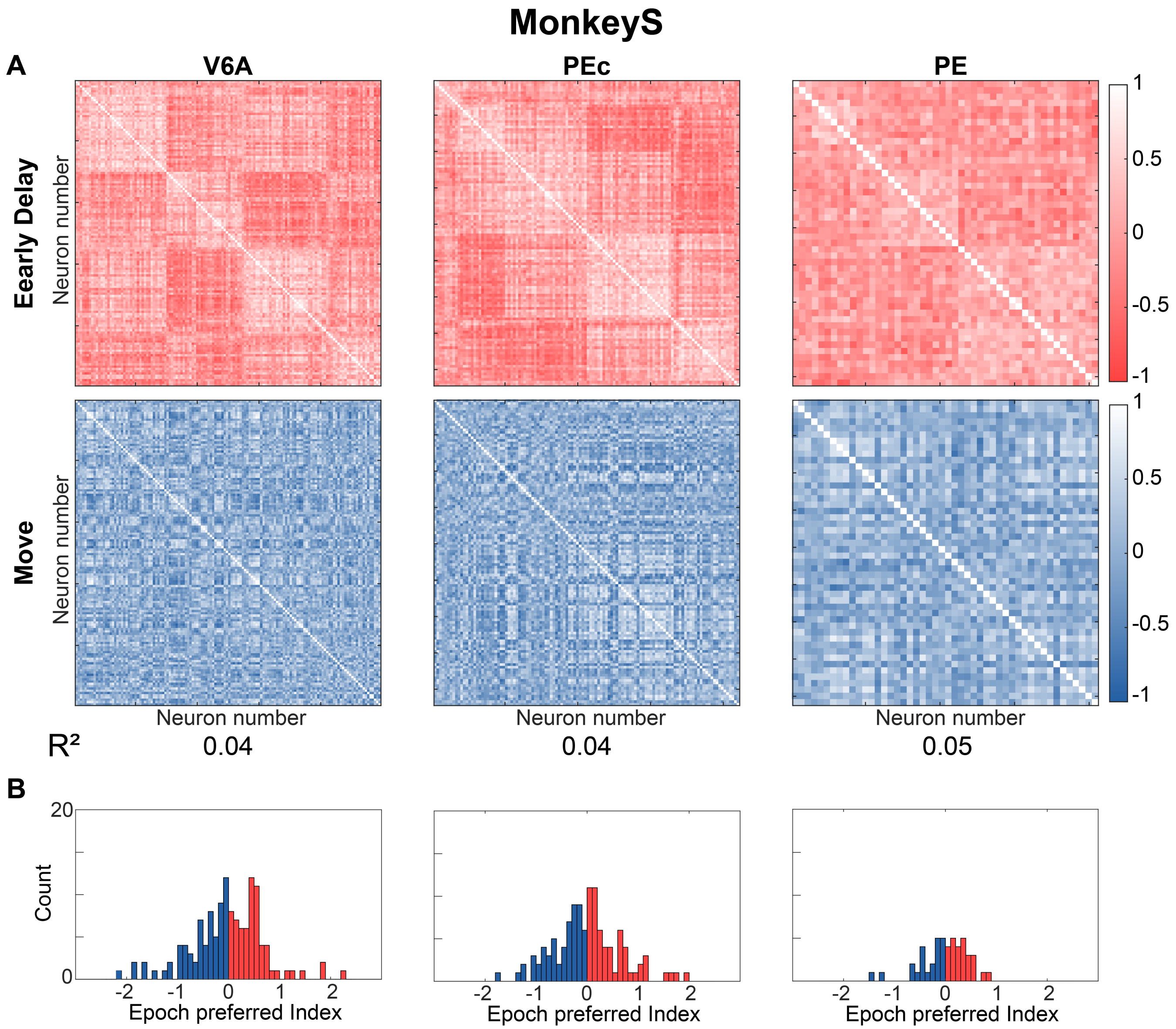


**Fig. S1. Pearson Cross-Correlation preliminary analysis for *MonkeyS***
Same format and analysis as in Figure 3, here applied to *MonkeyS*. **A**. Correlation matrices for the Early Delay epoch(Red) andMove epoch (Blue) epochs for all neurons in each area. The coefficient of determination $(R^2)$ quantifies how much of the correlation structure is preserved between the two behavioral periods. **B**. Distribution of the epoch-preference index across neurons (Hartigan's dip test, all p-val > 0.5)

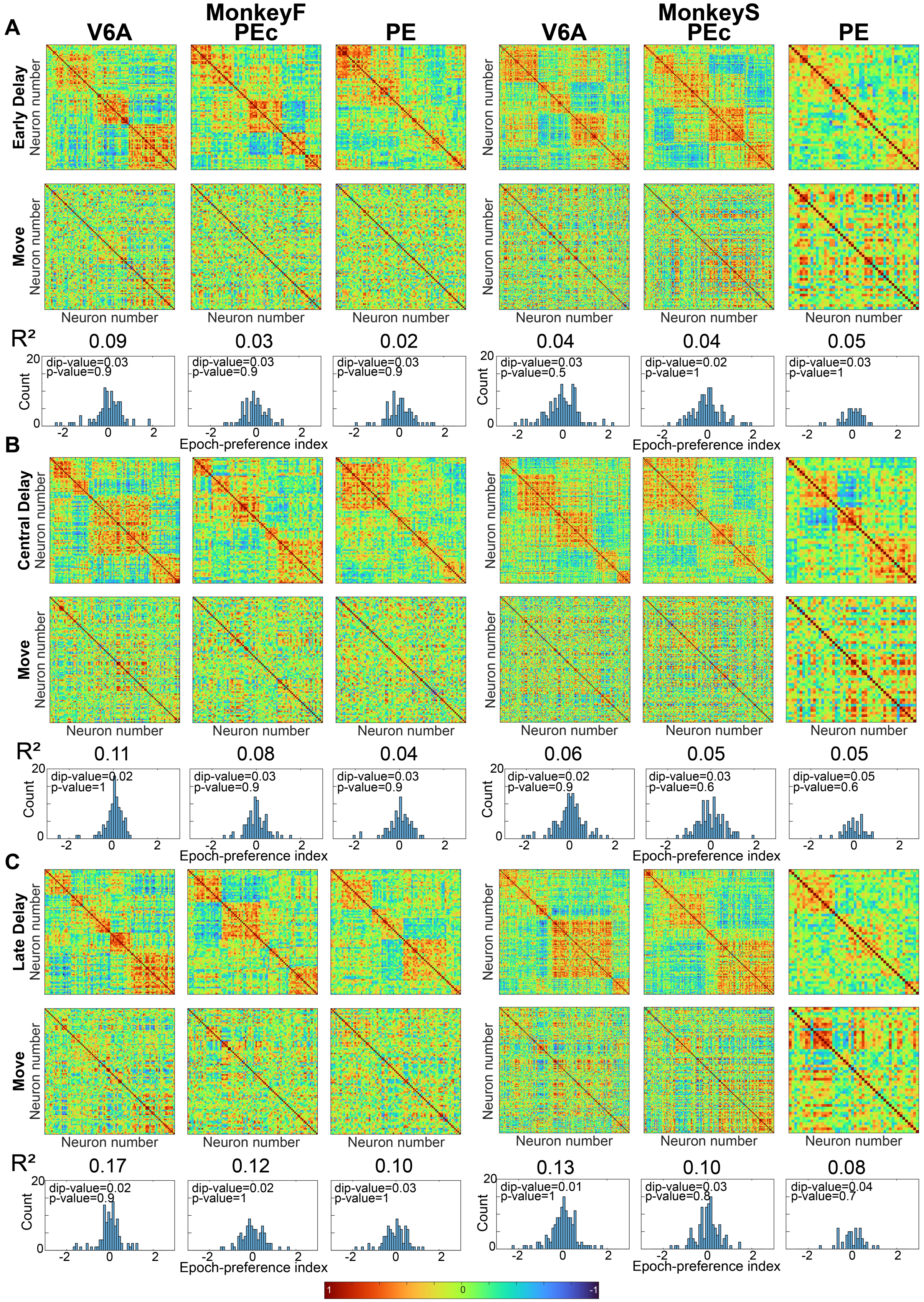


**Fig. S2. Pearson Cross-Correlation preliminary analysis across all epochs in *MonkeyS* and *MonkeyF***

For each animal (1-3 columns *MonkeyF* , 4-6 columns *MonkeyS*) and for each areas, each panel shows the comparison between one of the delay epochs and the epoch of movement: **A**. Early Delay vs Move epoch **B**. Central Delay vs Move epoch **C**.Late Delay vs Move epoch . In each histogram of the epoch-preference index we have reported the Hartigan's dip test results: dip statistic and p-value.

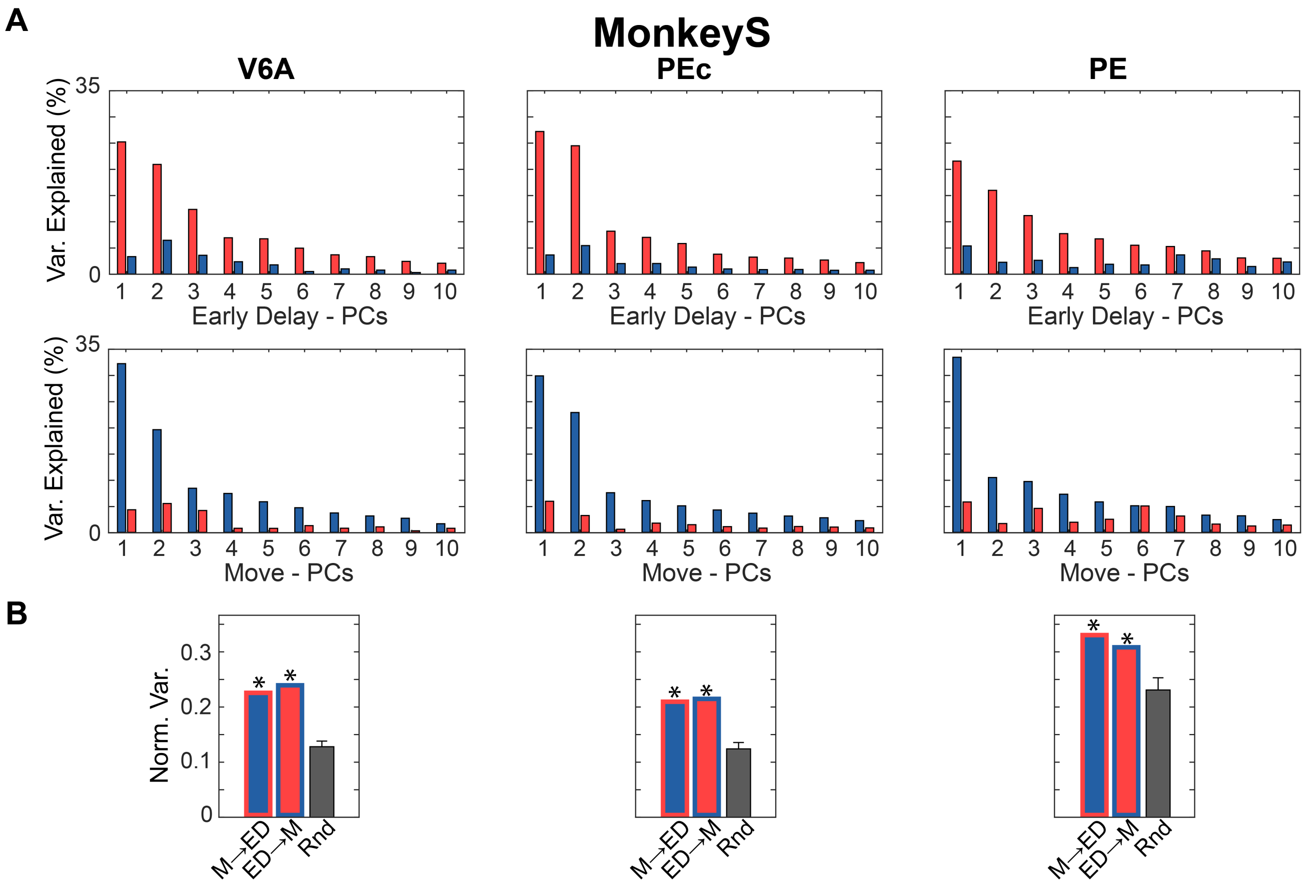


**Fig. S3. Classic Principal Component Analysis for *MonkeyS***
Same format and analysis as in Figure 3, here applied to *MonkeyS*. **A**.Variance explained in Early Delay (red) and Move (blue) epochs by principal components computed from each respective epoch. **B**. Normalized variance captured by cross-projections of neural activity onto 10-D subspaces. Bar colors and outlines follow the same conventions as in Figure 3. Asterisks indicate normalized cross variance significantly greater than chance (all p-val<0.001)

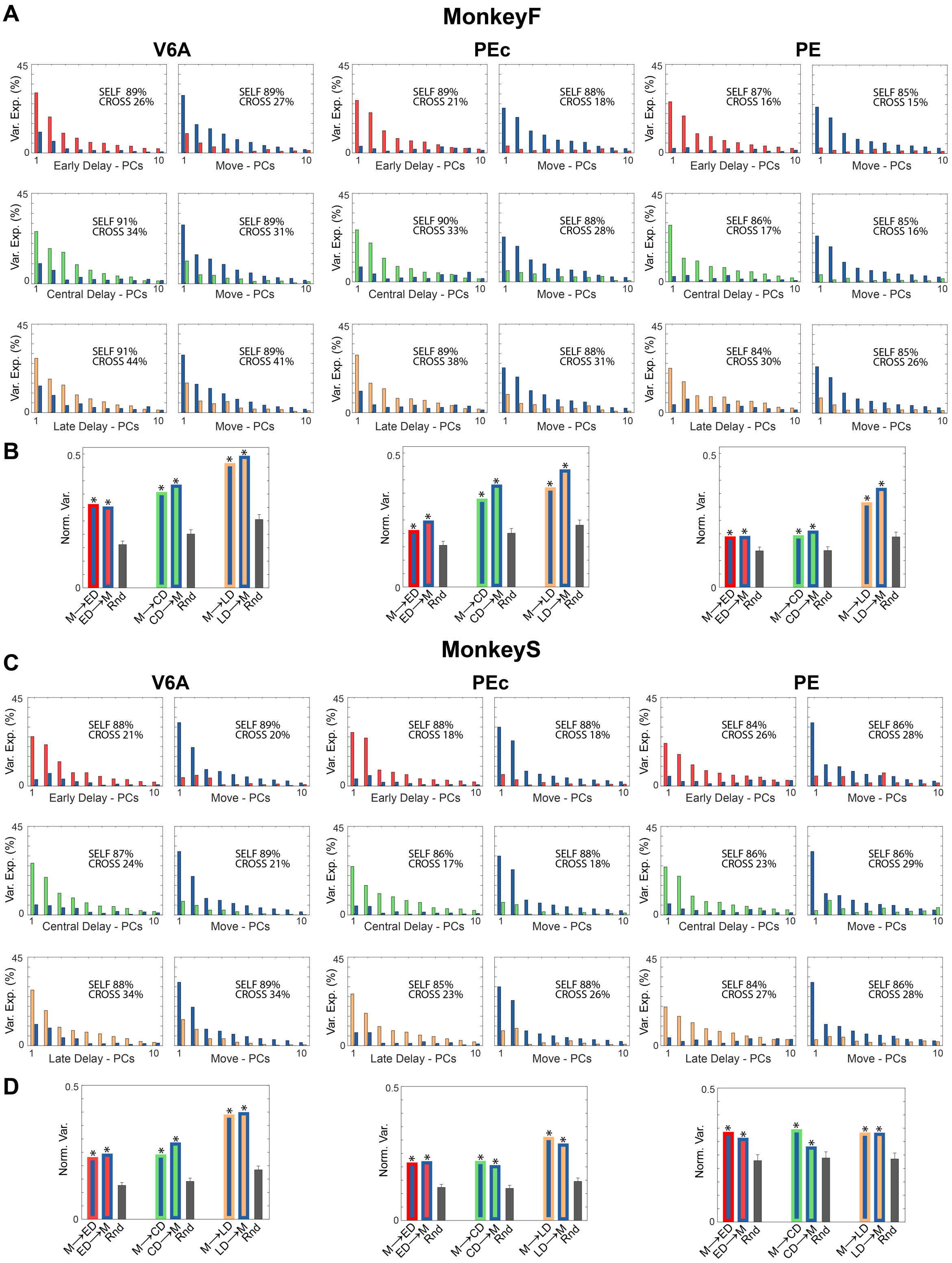


**Fig. S4. Classic Principal Component Analysis across all epochs in *MonkeyS* and *MonkeyF***
(**A**, **C**) Variance explained by the top 10 principal components (PCs) computed separately for Early Delay (ED, red), Central Delay (CD, green), Late Delay (LD, orange), and Movement (M, blue) epochs. Bars show variance explained when projecting activity from each epoch onto its own PCs (self-projection) and onto PCs from other epochs (cross-projection). Panel A shows data from Monkey F, Panel C from Monkey S. (**B**, **D**) Normalized cross-projected variance between all pairs of epochs. Bar fill color indicates the projected epoch; bar outline color indicates the subspace (PCs) used. Gray bars represent the chance level (mean ± SD across 10,000 random subspaces).Asterisks shows if the normalized cross variance is significantly higher than chance (p-val<0.05, one-tailed test). Panel B shows data from Monkey F, Panel D from Monkey S. Abbreviations: ED – Early Delay, CD – Central Delay, LD – Late Delay, M – Movement, Rnd – Random.

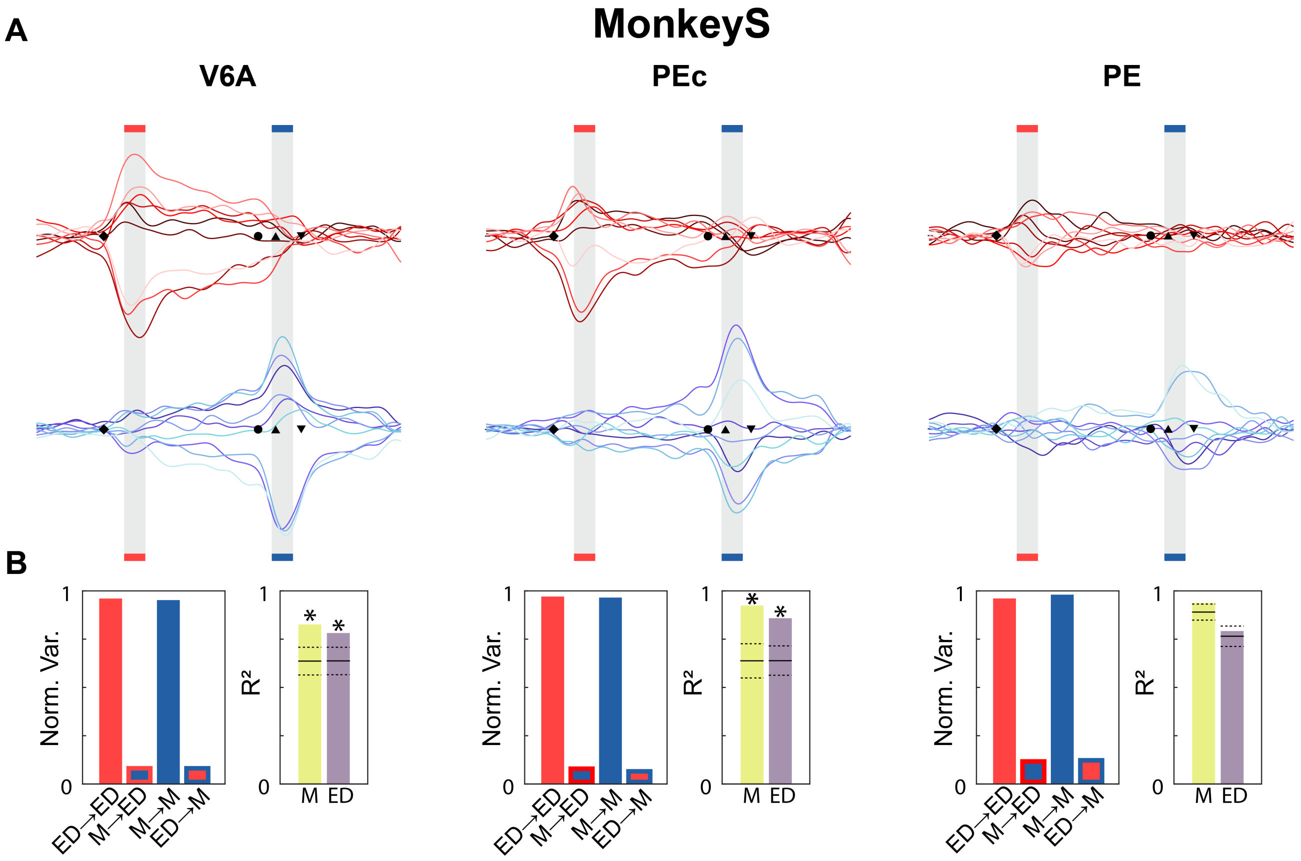


**Fig. S5. Orthogonal Subspaces analysis for *MonkeyS***

Same format and analysis as in Figure 4, here applied to *MonkeyS*. **A** Projections of neural population activity onto the first principal component of the Early Delay orthogonal subspace (red traces) and the Move orthogonal subspace (blue traces). Different color intensities represent different reach directions, from left (darkest) to right (lightest). **B** Two plots are shown per area. Left: Normalized variance of neural activity from each epoch projected onto each orthogonal subspace. Stroke color indicates the subspace used; fill color indicates the activity epoch. For example, a blue bar with red stroke (M→ED) shows variance of Move activity projected onto the Early Delay subspace; a red bar with red stroke (ED→ED) shows Early Delay activity projected onto its own subspace. Right: Coefficient of determination $(R^2)$ for linear estimations between dynamics across epochs. The yellow bar shows $R^2$ of estimating Move activity in the Move subspace using Early Delay projections in the Early Delay subspace; the pink bar shows the inverse (ED from M). Asterisks indicate statistically significant fits (p-val $<$ 0.05, one-tailed test). Null distributions were obtained by shuffling the independent variable 10,000 times; the mean and standard deviation are shown as solid and dashed horizontal lines, respectively. Abbreviations: ED – Early Delay, M – Move.

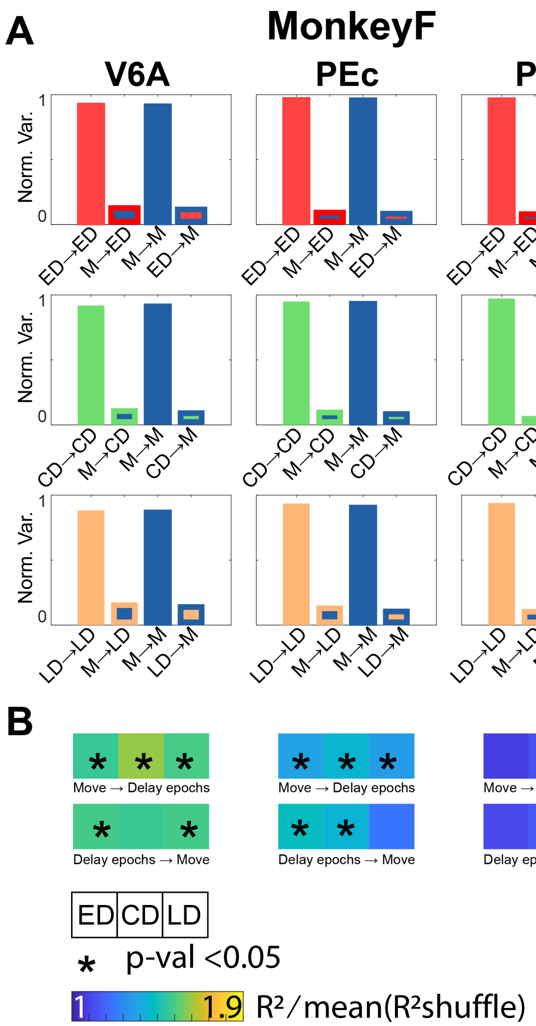


**Fig. S6. Orthogonal Subspaces analysis across all epochs in *MonkeyS* and *MonkeyF***
**A** For both animals and for all areas each graph shows the self and cross Normalized Variance in the Early/Central/Late Delay and Move subspaces. Color code as to be interpreted as a follow: the stroke color of the column represent the subspaces in which neural activity was projected, the fill color instead represent which neural activity was been projected. For instance Blue column whit red stroke represent the normalized variance of the Move activity projected on the Early Delay subspaces (M→ED), instead red column whit red stroke is the normalized variance of the Early Delay activity projected on the Early Delay subspaces (ED→ED). **B** Linear estimation result in terms of $R^2$. In the first row (Move → Delay epochs) Move epoch projected onto its subspaces has been use to estimate the Early/Central/Late Delay epoch projected onto their subspace. Second row is the opposite(Delay epochs → Move). Asterisks shows if the fitting is significantly different from a null distribution ($p - val < 0.05$, one-tailed test). Null distribution was obtained shuffling the independent variable of the fitting. This shuffled version of the fitting has been repeated $10000$ times and correspond to our null distribution. The mean of the distribution is represented by horizontal continuous line and the standard deviation with a horizontal dashed line. Colors represent the ratio between $R^2$ obtained by linear estimation and the mean $R^2$ of the null distribution. Abbreviations: ED, Early Delay; CD, Central Delay; LD, Late Delay; M, Move.

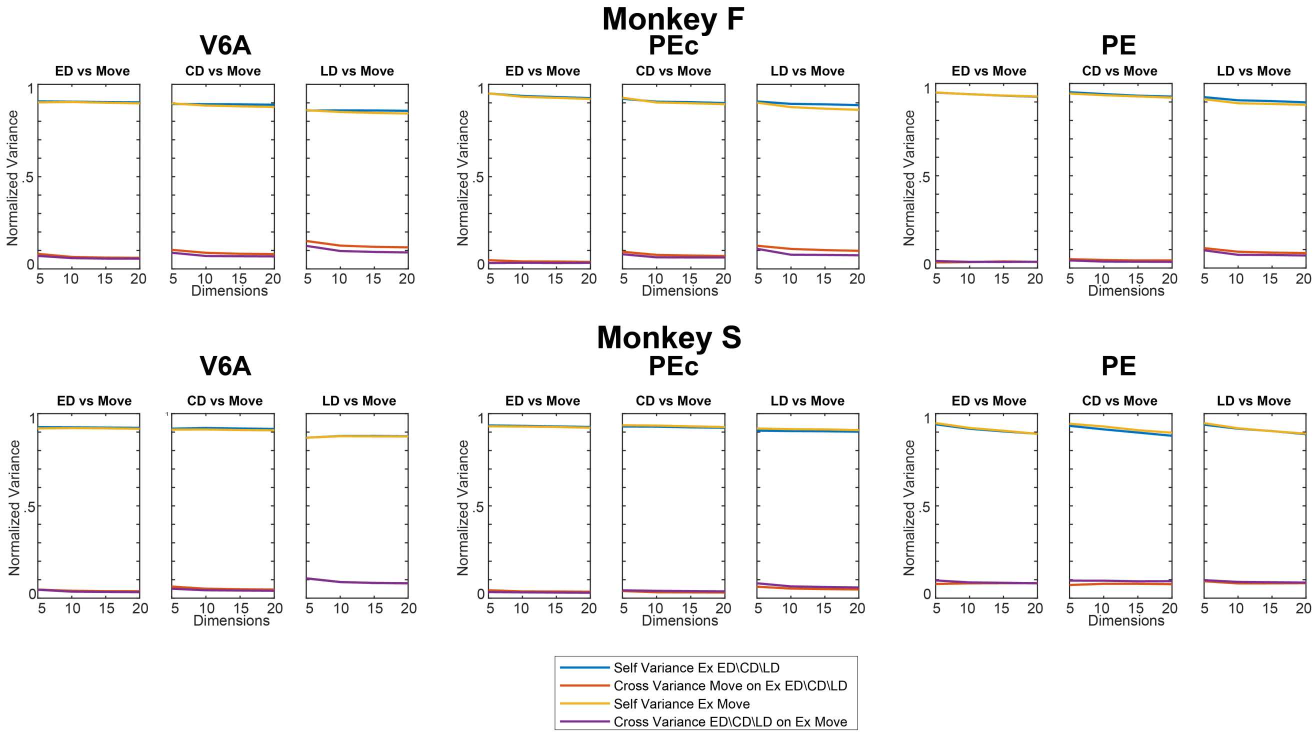


**Fig. S7. Multi-dimensional Orthogonal Subspaces analysis across all epochs in *MonkeyS* and *MonkeyF***
For both animals, the normalized variance with respect to the dimension considered for the orthogonal subspaces was calculated for all areas and epochs. Abbreviations: ED, Early Delay; CD, Central Delay; LD, Late Delay; M, Move.

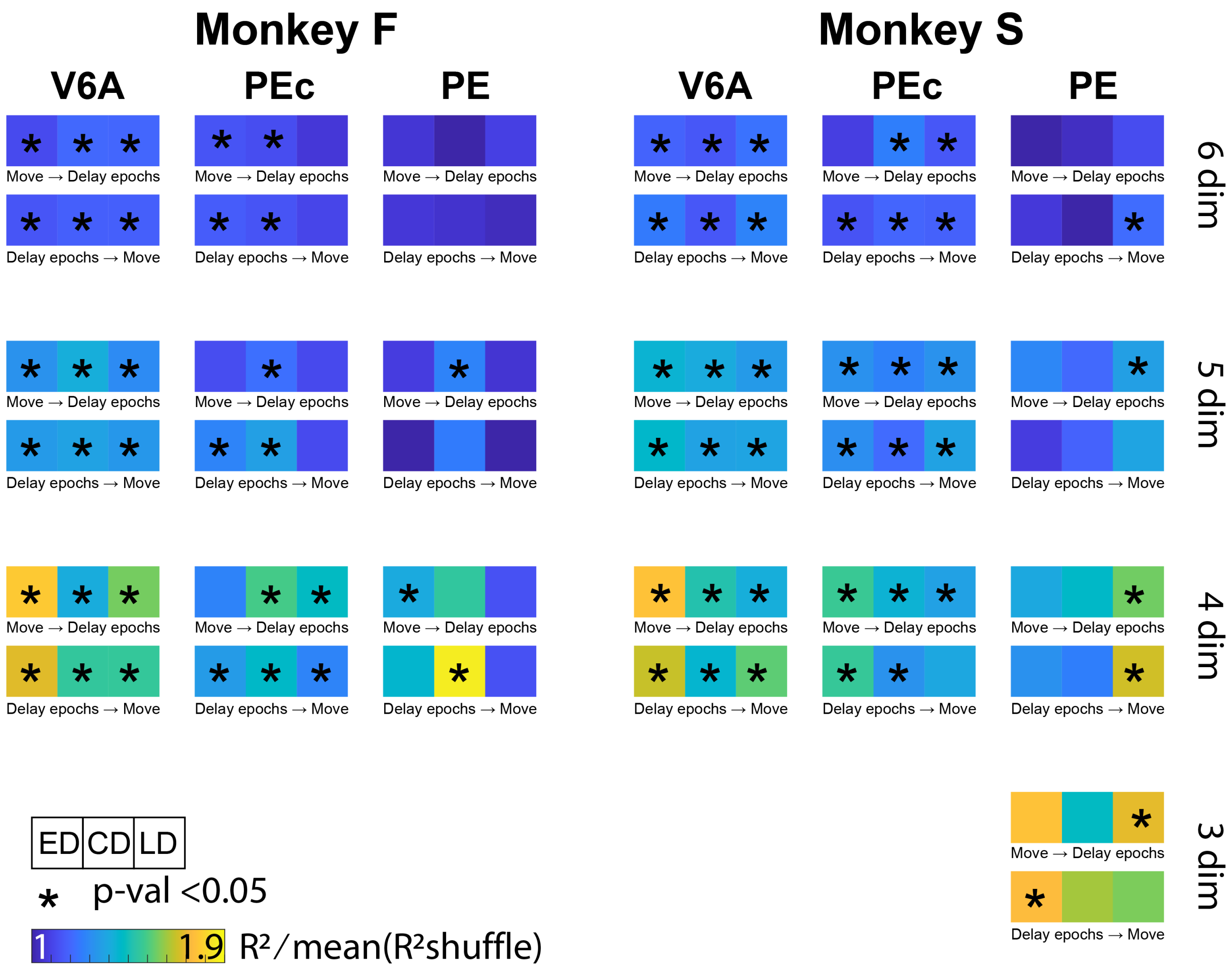


**Fig. S8. Multi-dimensional Linear Estimation analysis across all epochs in *MonkeyS* and *MonkeyF***
Linear estimation result in terms of $R^2$. in the first row (Move→Delay epochs) Move epoch projected onto its subspaces has been use to estimate the Early/Central/Late Delay epoch projected onto their subspace. Second row is the opposite (Delay epochs→Move). Asterisks shows if the fitting is significantly different from a null distribution (p-val<0.05, one-tailed test). Null distribution was obtained shuffling the independent variable of the fitting. This shuffled version of the fitting has been repeated 10000 times and correspond to our null distribution. The mean of the distribution is represented by horizontal continuous line and the standard deviation with a horizontal dashed line. Colors represent the ratio between $R^2$ obtained by linear estimation and the mean $R^2$ of the null distribution. In each row analysis has been reported respect to the dimension of the subspace. Abbreviations: ED, Early Delay; CD, Central Delay; LD, Late Delay; M, Move.

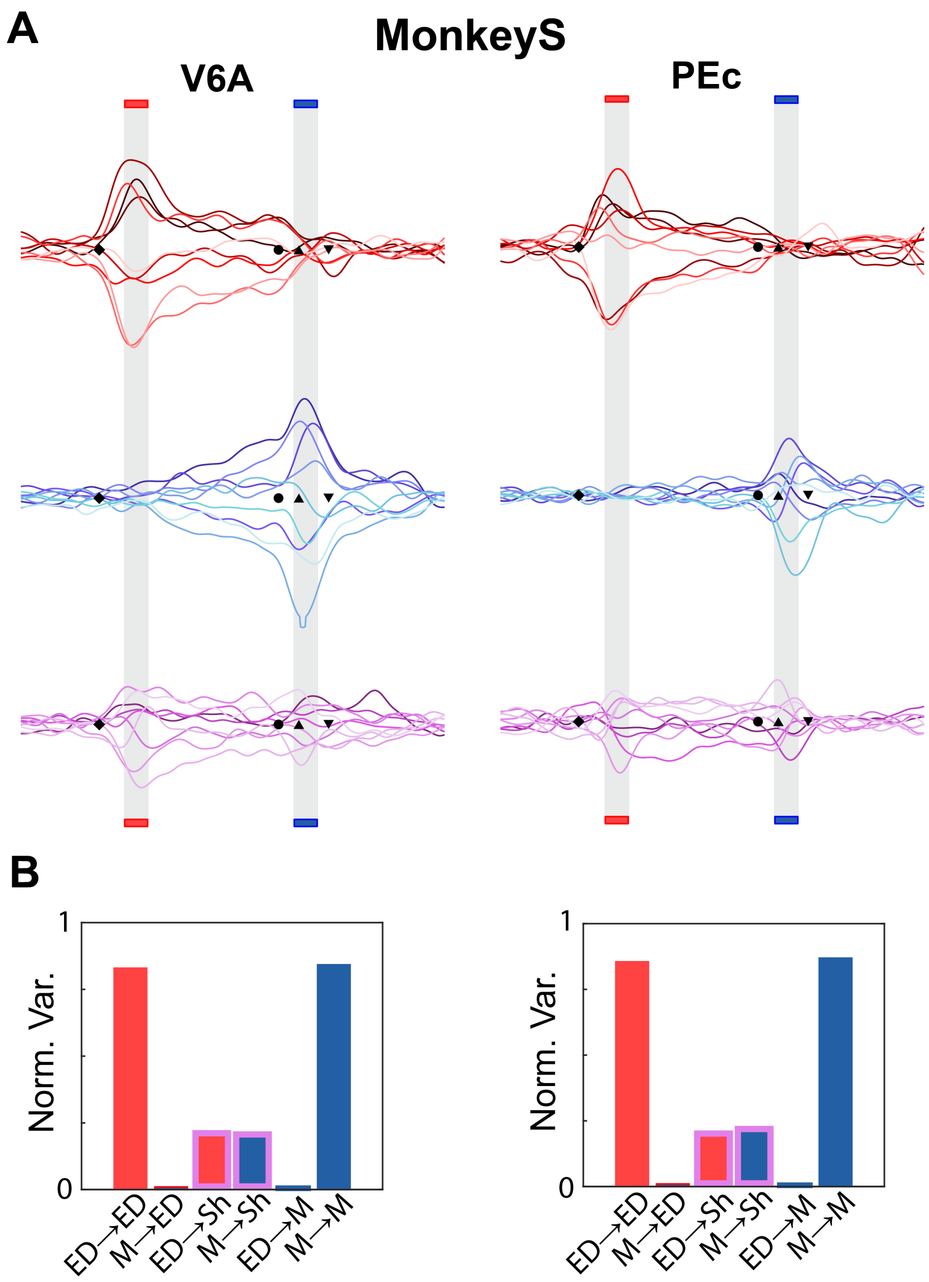


**Fig. S9. Exclusive and Shared subspaces analysis for *MonkeyS***
Same format and analysis as in Figure 5, here applied to *MonkeyS*. **A** Projections of the neural population activity on the first principal component.**B** Histogram shows the Normalized Variance relative to Exclusive and Shared subspaces. Abbreviations: ED, Early Delay; M, Move; Sh, Shared.

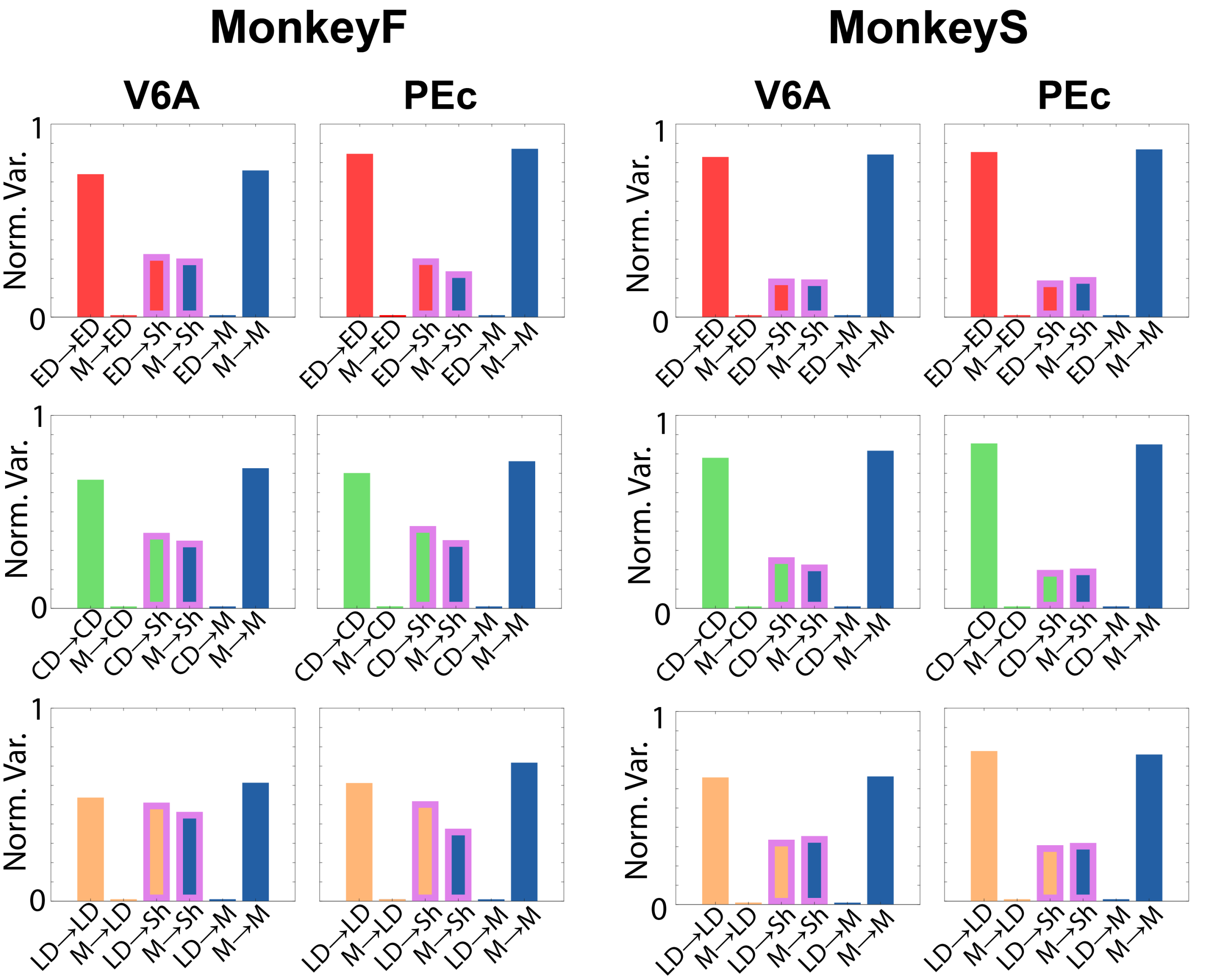


**Fig. S10. Exclusive and Shared subspaces analysis across all epochs in *MonkeyS* and *MonkeyF***
Histogram shows the Normalized Variance relative to Exclusive and Shared subspaces: the stroke color of the column represent the subspaces in which neural activity was projected (red, green, orange, blue for Early/Central/Late Delay and Move exclusive subspaces and magenta for Shared), the fill color instead represent which neural activity was been projected (ED red and M blue). For instance Blue column whit red stroke represent the normalized variance of the move activity projected on the Early Delay Exclusive subspaces (M→ED), instead red column whit magenta stroke is the normalized variance of the preparatory activity projected on the Shared subspaces (ED→Sh). Note that in the case of M→ED and ED→M only the stroke color of the bar is visible due to the low variance explained ($\sim 1\%$). Abbreviations: ED, Early Delay; CD, Central Delay; LD Late Date; M, Move; Sh, Shared.

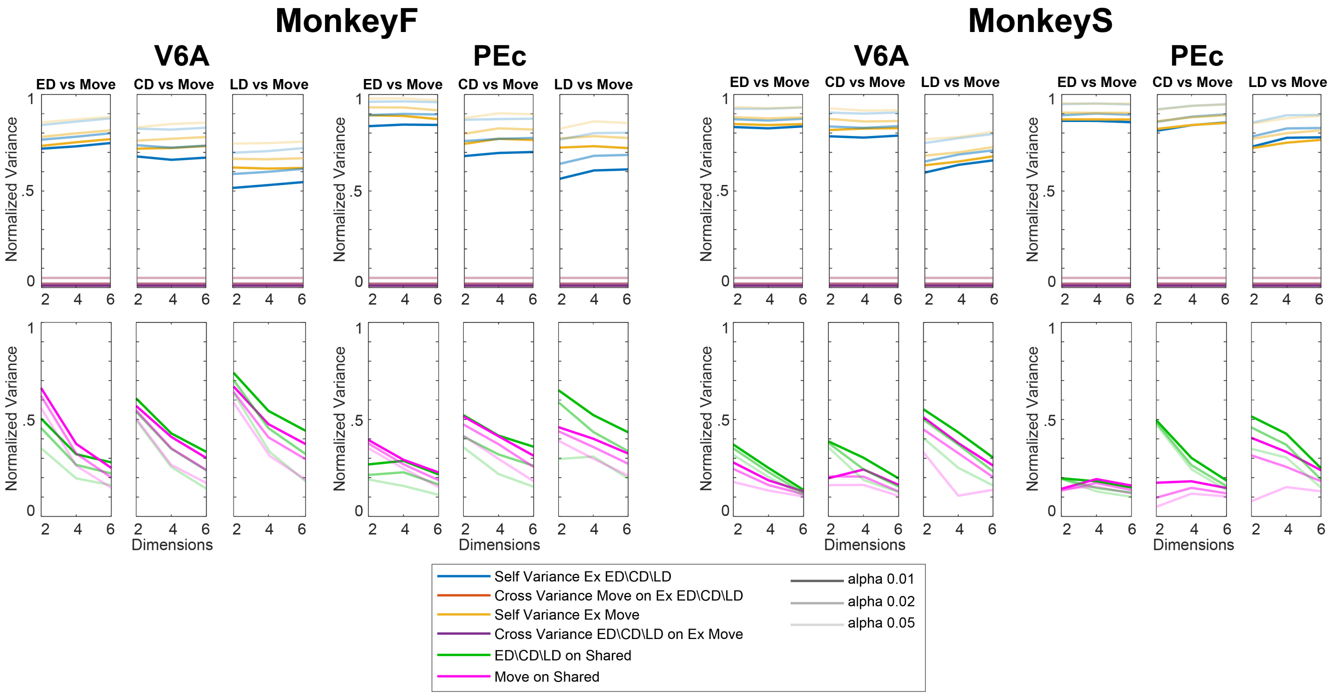


**Fig. S11. Multi-dimensional Exclusive and Shared subspaces analysis across all epochs in *MonkeyS* and *MonkeyF***

For both animals, the normalized variance with respect to the dimension considered for the Exclusive and Shared subspaces was calculated for all areas and epochs. Abbreviations: ED, Early Delay; CD, Central Delay; LD, Late Delay; M, Move.
